# Deconstructing hunting behavior reveals a tightly coupled stimulus-response loop

**DOI:** 10.1101/656959

**Authors:** Duncan S. Mearns, Julia L. Semmelhack, Joseph C. Donovan, Herwig Baier

## Abstract

Animals build behavioral sequences out of simple stereotyped actions. A comprehensive characterization of these actions and the rules underlying their temporal organization is necessary to understand sensorimotor transformations performed by the brain. Here, we use unsupervised methods to study behavioral sequences in zebrafish larvae. Generating a map of swim bouts, we reveal that fish modulate their tail movements along a continuum. We cluster bouts that share common kinematic features and contribute to similar behavioral sequences into seven modules. Behavioral sequences comprising a subset of modules bring prey into the anterior dorsal visual field of the larvae. Fish then release a capture maneuver comprising a stereotyped jaw movement and fine-tuned stereotyped tail movements to capture prey at various distances. We demonstrate that changes to chaining dynamics, but not module production, underlie prey capture deficits in two visually impaired mutants. Our analysis thus reveals the temporal organization of a vertebrate hunting behavior, with the implication that different neural architectures underlie prey pursuit and capture.

## Introduction

Quantitative descriptions of behavior are essential if we are to fully understand the brain (Krakauer et al., 2017). Such descriptions have provided a framework for interrogating the genetic and neural basis of behavior in worms, flies and mice (Cande et al., 2018; Kato et al., 2015; Vogelstein et al., 2014; Wiltschko et al., 2015). It is believed that complex, flexible behavior arises as a result of animals chaining together simpler, more stereotyped movements (Anderson and Perona, 2014; Egnor and Branson, 2016; Tinbergen, 1951). These movements can be generated spontaneously through internal neural processes and/or induced by external stimuli impinging on the animal’s sensory organs. Thus, a comprehensive model of an animal’s behavior should identify the constituent building blocks of the behavior, uncover rules governing the chaining of these building blocks into longer sequences, and account for how the animal’s sensory experience shapes and guides these sequences (Coen et al., 2014; Seeds et al., 2014; Tinbergen, 1951; Wiltschko et al., 2015). Such an account of behavior could uncover the sensorimotor transformations performed by the brain that are critical for survival in a dynamically changing world.

The individual movement patterns that constitute behavior have been termed motor primitives (Flash and Hochner, 2005), synergies (Bizzi and Cheung, 2013), movemes (Del Vecchio et al., 2003), or behavioral modules (Berman et al., 2014; Brown et al., 2013; Egnor and Branson, 2016; Marques et al., 2018; Wiltschko et al., 2015). However, whether such modules truly constitute stereotyped, invariant movements or whether they merely reflect extremes in a behavioral continuum remains unclear (Berman et al., 2014; Katsov et al., 2017; Marques et al., 2018; Patterson et al., 2013; Szigeti et al., 2015). In either case, actions must be chained into sequences that reliably achieve the desired goal of the animal. Stereotyped, reproducible behavioral sequences have been explained with serial models, in which one action triggers the next in the chain via feed-forward neural mechanisms (Long et al., 2010). In contrast, flexible sequences, in which the ordering of modules might be different each time the behavior occurs, have been explained using hierarchical models. In hierarchical models, switching between behavioral modules is stochastic, but may be influenced by longer-term behavioral states or sensory stimuli received by the animal (Berman et al., 2016; Seeds et al., 2014; Tao et al., 2019; Wiltschko et al., 2015).

Capturing prey is an essential behavior for the survival of many animals and is innate. The behavior is also complex, requiring the localization, pursuit and capture of a prey object, often moving in a three-dimensional environment. Consequently the action sequences that constitute this behavior are required to be flexible, allowing animals to adapt to the specific movement of the current stimulus (Ewert, 1987). Zebrafish larvae hunt protists that float in the water column (Borla et al., 2002; Budick and O’Malley, 2000; McElligott and O’Malley, 2005). Larvae do not perform continuous locomotion, but rather swim in discrete bouts with a beat-and-glide structure (Budick and O’Malley, 2000), which aids the segmentation of their behavior into discrete actions (Marques et al., 2018). Both real and virtual prey presented to restrained animals can produce isolated orienting swim bouts and eye convergence: hallmarks of prey capture in zebrafish larvae (Bianco et al., 2011; Semmelhack et al., 2014). It has been suggested that such movements could compound over time in a stimulus-response loop, whereby movements of the tail and eyes bring prey to the near-anterior visual field of the animals (Patterson et al., 2013; Trivedi and Bollmann, 2013). However, it is not clear whether such a model would be implemented by gradual changes in the kinematics of bouts over the course of a hunting sequence (Borla et al., 2002; Patterson et al., 2013), or as a result of discrete switches between more stereotyped motor patterns (Marques et al., 2018). One possibility, that has not been tested, is that different stages of the behavior have a different organization. For example, animals might dynamically modulate their movements to adjust to the position of the prey during pursuit, but resort to more stereotyped action patterns when consuming prey (Ewert, 1987). Moreover, studies of prey capture have predominantly focused only on either tail, jaw, or fin movements and it is not known how these movements are coordinated into temporally organized patterns over the entire behavioral sequence (Borla et al., 2002; Hernández et al., 2002; McClenahan et al., 2012; Patterson et al., 2013).

Here, we present a novel computational framework for generating a map of movements made by an animal. We apply our algorithm to the bouts of week-old zebrafish larvae swimming in the presence of prey and find a continuum of behaviors. In this continuous space we identify seven modules, which correspond to groups of bouts with similar kinematics and that also share common transitions to and from other modules. Sequences of bouts through a subset of these modules are reproducible across prey capture events due to a tightly coupled stimulus-response loop, in which the fish’s movements generate new stimuli that trigger subsequent bouts in the chain. Further investigating the capture strike, during which prey are consumed, we show that variation in this behavior arises from differential chaining of stereotyped tail and jaw movements, mediated by prey distance. We validate our behavioral classification by showing genetic differences in the initiation and chaining of prey capture modules in mutants with impaired visual processing.

## Results

### Swim bouts are trajectories through a low-dimensional postural space

To study the organization of prey capture in zebrafish larvae, we first sought to characterize the basic building blocks of this behavior. To this end, we recorded individual larvae (7-8 dpf; n=45; 20 min each) hunting live prey (paramecia) in a custom-built behavioral arena (**Figure 1A**) and tracked the tail and eyes of the fish in each frame (**Figure 1B,C** and **Video 1**; see Methods). From this dataset, we automatically identified and segmented 57,644 individual swim bouts for future analysis.

**Figure 1.**
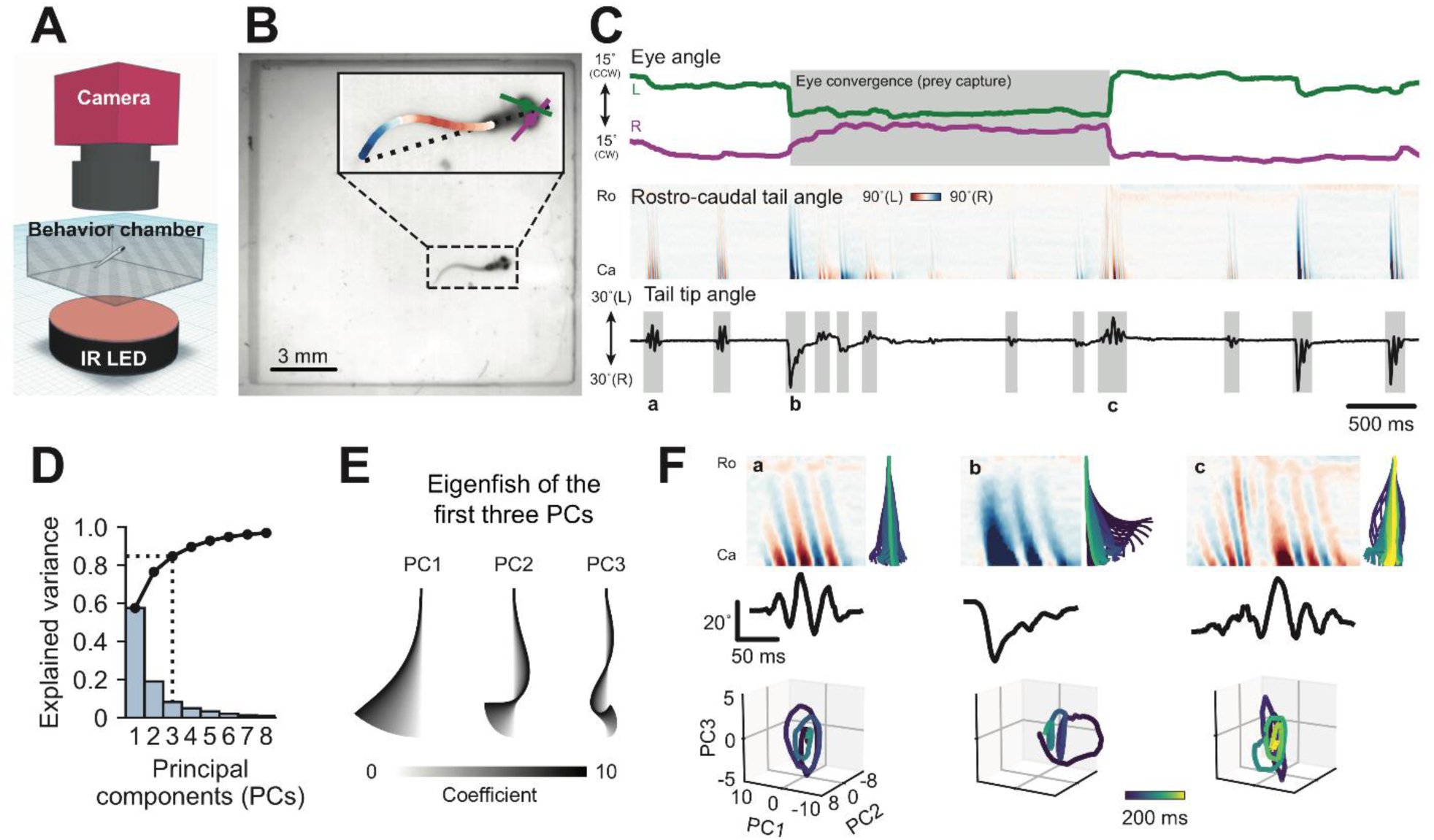
Swim bouts are characterized by their postural dynamics. (**A**) Schematic of the setup used to record behavioral data. (**B**) Example frame from high-speed video recording. Inset is overlaid with tail and eye tracking. (**C**) Eye and tail kinematics extracted from six seconds of behavioral recording. Ro: Rostral, Ca: Caudal. (**D**) Principal component analysis of tail shapes. Explained variance (bars) and cumulative explained variance (points) of the first eight components. We retained three components, which explained 84.7% of the variance. (**E**) Schematized “eigenfish” of the first three principal components. (**F**) Individual bouts represented by trajectories through postural space. Top left panels: curvature along rostral-caudal axis of the tail over time. Bottom panels: bouts represented by trajectories through the first three principal components. Top right panels: tail movement reconstructed from these trajectories.

To reduce the dimensionality of this vast dataset, we applied principal component analysis (PCA) to the tail kinematics of the fish during swim bouts and found that just three components were sufficient to explain 84.7% of the variance in tail posture (**Figure 1D**). These principal components define postural modes and can be represented by a set of “eigenfish” (Girdhar et al., 2015; Stephens et al., 2008; Szigeti et al., 2015), which show the unmixed tail shapes encoded by each component (**Figure 1E**). As the posture of the animal evolves over time, the changing tail shape traces a trajectory in the three-dimensional coordinate space defined by the postural modes (**Figure 1F** and **Video 2**). Thus we find that the tail kinematics of zebrafish larvae are inherently low-dimensional, which provides a useful way to represent bouts for subsequent analysis.

### Task-specific motor programs occupy distinct domains of the behavioral space

Next, we wanted to know whether animals build their behavioral sequences from kinematically discrete motor programs or draw their bouts from a continuous behavioral manifold. We sought to distinguish these possibilities by representing swim bouts in a space where neighboring points encode bouts with similar postural dynamics. In this space, tight clusters would suggest that larvae can only generate a limited number of stereotyped bout types, whereas a diffuse cloud would suggest that larvae can continuously modulate the kinematics of their bouts. To distinguish these possibilities, we developed a pipeline for determining the structure of the behavioral manifold (**Figure 2A**; see Methods). Our algorithm consists of three steps: alignment, clustering and embedding. In the first step, we calculate the distance between each pair of bouts in the three-dimensional postural space using dynamic time warping (DTW) (Jouary and Sumbre, 2016; Sakoe and Chiba, 1978). Next, we performed a round of affinity propagation (Frey and Dueck, 2007) prior to embedding, using the negative DTW distance between a given pair of bouts as a measure of their similarity. Using the median similarity between bouts as the basis for affinity propagation produced 1,744 clusters containing at least three bouts. Since affinity propagation identifies an exemplar to represent each cluster, we produced our final behavioral space by performing isomap embedding (Tenenbaum et al., 2000) of these exemplars. For the isomap embedding, we constructed a nearest-neighbors graph of the exemplars using their DTW distances, and calculated the minimum distance between each pair of points in this graph. We used three dimensions for this final behavioral space to minimize the reconstruction error of the embedding with as few dimensions as possible (**Figure 2 – figure supplement 1A**), as well as to maximize the interpretability of bout separation in the resulting space.

**Figure 2.**
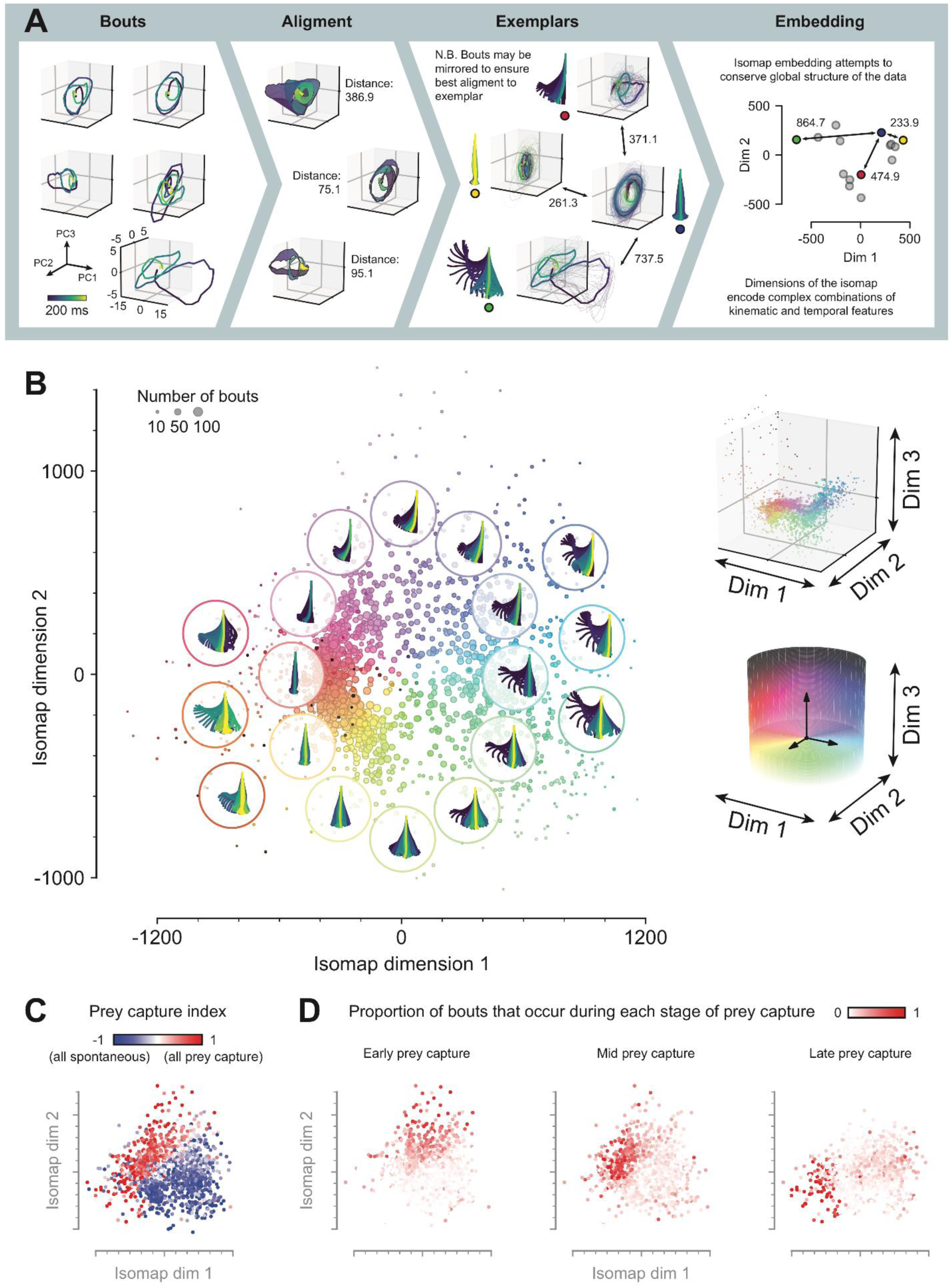
Generation of a zebrafish larva behavioral space. (**A**) Analysis pipeline for generating behavioral space. Each bout is a trajectory through a three dimensional postural space. Pairwise distances between all bouts are computed using dynamic time warping (DTW). Nearby bouts are grouped and a representative exemplar is chosen from each small group. Exemplars are then embedded in a low-dimensional behavioral space using isomap embedding on their DTW distances. (**B**) Behavioral space. Left: representative bouts projected onto the first two dimensions of the behavioral space. Right: behavioral space rendered in all three dimensions. Points colored according to position within a hue-lightness cylinder centered on origin. (**C**) Prey capture index (defined using eye convergence) of each exemplar in the behavioral space. Index defined as (# prey capture bouts - # spontaneous bouts) / (# total bouts) mapped to each exemplar. (**D**) Proportion of bouts mapped to each exemplar that occur during early, middle or late phases of prey capture. See also **Figure 2 – figure supplement 1**,**2**.

Inspecting this behavioral space, we do not observe tight clusters with stereotyped kinematic features, but rather loosely clustered bout types separated by more sparsely populated regions (**Figure 2B** and **Figure 2 – figure supplement 1B**). Such a structure suggests a behavioral continuum, with certain motifs favored in our particular behavioral paradigm. Inspecting bouts that are represented in different regions of the space, we find high-amplitude bouts with a late turning component (far left, dark warm color), forward scoots (lower left, red to green colors), turns (right, green to blue colors), and asymmetric bouts (top, purple to magenta colors). This suggests that turn angle and swimming speed are the dominant kinematic features that define larval swim bouts (**Figure 2 – figure supplement 1C**). Turn angle and angular velocity separate bouts along the first dimension of the space, and swimming speed separates bouts along the second and third dimensions.

Next, we sought to determine where prey capture bouts lie in the behavioral space, and to what extent they are kinematically distinct from spontaneous swims. To this end, we used eye convergence as an independent and unbiased indicator of prey capture behavior (Bianco et al., 2011) (**Figure 2 – figure supplement 2**). We assigned each point in the space a prey capture index, indicating how frequently each bout was recruited during prey capture (eyes converged) versus spontaneous behaviors (eyes not converged). Markedly, we found that prey capture and spontaneous bouts were clearly separated in the behavioral space (**Figure 2C**). Furthermore, when we decomposed prey capture swims into early, mid and late bouts of a hunting sequence, we found further delineation in the behavioral space (**Figure 2D**). These results reveal that distinct motor programs are differentially recruited during hunting and spontaneous swimming and that larvae systematically alter the kinematics of their bouts over the course of a hunting sequence.

### Behavioral sequences are built from a small number of simple transition modes

Having identified the kinematic structure of zebrafish larva swim bouts, we next wanted to investigate how the temporal organization of bouts produced behavioral sequences (**Figure 3A**). We reasoned that, despite the large number of bouts that populate the behavioral space, the goal-oriented nature of prey capture behavior would produce stereotyped sequences through this space. To test this possibility, we generated a transition frequency matrix from the number of transitions between each cluster in behavioral space. To distinguish between symmetric transitions, where animals stay in the same part of the behavioral space (i.e. repeating bouts with shared kinematic features), and asymmetric transitions, where animals transition to a different part of the space (i.e. switching to a different kind of behavior), we decomposed the matrix into its symmetric and antisymmetric parts. We then factorized the symmetric and antisymmetric matrices using singular-value decomposition (SVD) to obtain symmetric and antisymmetric transition modes (**Figure 3 – figure supplement 1**; see Methods). A symmetric transition mode describes transitions within a region of the behavioral space. Transitions from one region of the space to another can be described by an antisymmetric transition mode: groups of bouts occupying different areas of the behavioral space that tend to transition in one direction preferentially over the other. Each transition mode is associated with a singular value, which describes the extent to which the mode contributes to all the transitions recorded in the data.

**Figure 3.**
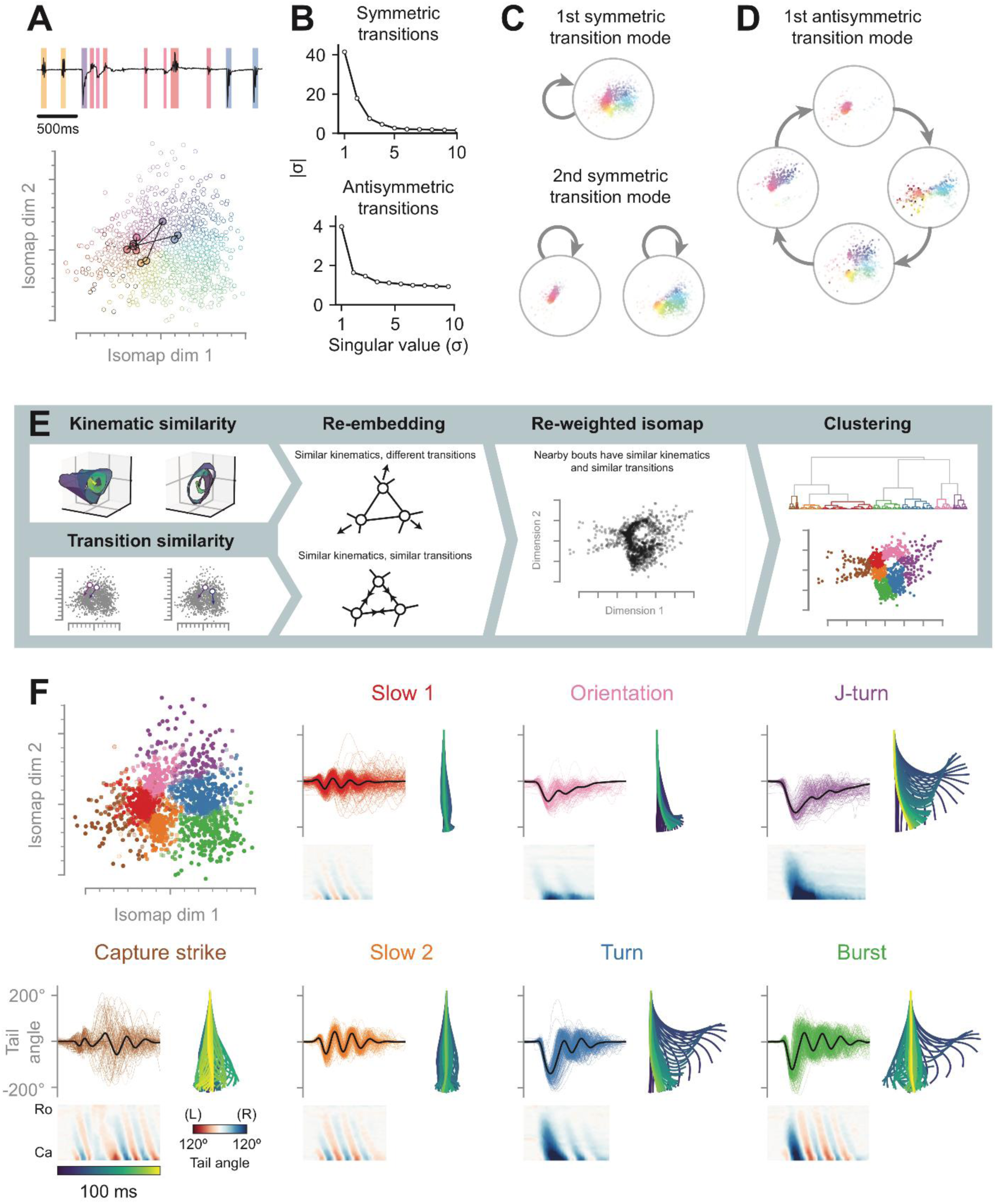
Behavior is composed of seven modules with distinct kinematics and dynamics. (**A**) Bouts are chained into sequences. Top: tail tip angle trace from **Figure 1C** with bouts color-coded according to position in the behavioral space. Bottom: same sequence plotted as a sequence through the behavioral space. (**B**) Singular-value decomposition (SVD) of the transition frequency matrix (from all observed transitions between bout pairs). The transition frequency matrix was smoothed and decomposed into its symmetric and antisymmetric components to identify transitions that occur in both directions and those that predominantly occur in only one direction (see **Figure 3 – figure supplement 1**; Methods). Top: singular values of the symmetric component of the transition matrix. Bottom: singular values of the antisymmetric component of the transition matrix. (**C**) First two symmetric transition modes. Symmetric transition modes describe transitions that occur within a group of bouts. (**D**) First antisymmetric transition mode. An antisymmetric transition mode encodes cyclic transitions through different groups of bouts. (**E**) Pipeline for generating a hybrid kinematic-transition space using re-weighted isomap embedding. Distances between exemplars in the behavioral space are rescaled by the distances between exemplars in the transition space defined by the transition modes. Bouts in the new space are clustered using hierarchical clustering. (**F**) Seven behavioral modules identified by hierarchical clustering in the kinematic-transition space. Top left: exemplars in the original behavioral space colored according to module. Subpanels show: individual tail angle traces in color with the average in black (top left); tail kinematics of a representative bout (bottom); tail reconstruction of the representative bout (right). See also **Figure 3 – figure supplement 1**,**2**.

Despite there being more than 3 million possible transitions between points in the behavioral space, we found that two symmetric and one antisymmetric transition mode accounted for most of the transitions in the data (**Figure 3B**, elbow in the singular values). Symmetric modes are represented by a single vector and antisymmetric modes by a pair of vectors; and each cluster in the behavioral space contributes either a positive or negative weight to each of these vectors (**Figure 3 – figure supplement 1B**). Therefore, to visualize which transitions were represented by each mode, we mapped these weights back into the behavioral space (**Figure 3C,D**). The first symmetric transition mode necessarily reflects the overall distribution of bouts in the behavioral space, since a majority of transitions occur between the most common types of bouts. The second symmetric mode separates low amplitude prey capture swims from spontaneous swims (**Figure 3C**). This indicates that animals often chain multiple low amplitude swims together, are less likely to chain low amplitude swims into spontaneous swims, and likewise less likely to chain spontaneous swims into low amplitude swims. The first antisymmetric transition mode represents transitions through different prey capture regions of the behavioral space (**Figure 2D** and **Figure 3D**). This suggests that transitions between different regions of the behavioral space tend to follow the sequence: asymmetric turn, low amplitude swim, which is then followed by either a “late prey capture swim” or spontaneous turn. In conclusion, we find different behavioral dynamics during self-generated spontaneous swimming and goal-oriented prey capture sequences in the zebrafish larva. Spontaneous swimming contains transitions between forward swims and turns that do appear to follow a specific sequence. On the other hands, prey capture sequences appear to be more structured, with bout kinematics systematically altered in a similar way over the course of the behavior each time it occurs.

### Bouts are organized into modules that tile the behavioral space

Bouts for exploratory and prey capture behavior form a continuum, and transitions between different regions of the behavioral space are explained by few transition modes. This suggested to us that behavior might be organized into modules, where each module represents a cluster of bouts with similar kinematics as well as similar transitions to and from other modules. Therefore, we generated a new kinematic-transition space, which contained information about both bout kinematics (from our behavioral embedding) and chaining structure (from our transition modes). We rescaled the graph distance between exemplars obtained using DTW by the corresponding distance between exemplars in a Euclidean space defined by transition modes; and proceeded with isomap embedding using this graph (**Figure 3E**; see Methods). Hierarchical clustering revealed seven modules that tile the original behavioral space (**Figure 3F**), many of which correspond to previously described bout types (Marques et al., 2018; McElligott and O’Malley, 2005; Patterson et al., 2013). We call these modules J-turns, orientations, “slow 1” swims, capture strikes, “slow 2” swims, burst swims and routine turns. J-turns, orientations, “slow 1” swims and capture strikes predominantly occurred during periods of eye convergence, and thus we term them prey capture modules (**Figure 3 – figure supplement 2**). In addition to these prey capture modules, we also identified three spontaneous swimming modules, “slow 2” swims, routine turns and burst swims, which predominantly occurred when the eyes were not converged. Thus, we find that despite the close juxtaposition of motifs in our behavioral space, nonetheless zebrafish larvae specifically recruit bouts from different regions of this space for different behavioral tasks. These regions correspond to behavioral modules that are not only kinematically distinct, but also occupy different positions within a behavioral chain.

### Prey capture sequences follow non-random, short-memory transition rules

Next, we investigated the temporal organization of prey capture and spontaneous swimming. On the one hand, behavior could be organized hierarchically, with animals switching between swimming states during which they preferentially perform bouts from only a subset of modules. Alternatively, animals could generate stereotyped sequences through modules, with individual modules shared between multiple sequences. To distinguish these possibilities, we constructed a family of models with different levels of memory about past behavior and tested the efficacy of these models in predicting the next bout in behavioral sequences (**Figure 4A,B**). In the first model, larvae randomly transitioned between bouts, with no impact from previous ones. This “memoryless” model provided a baseline performance against which other models could be compared. Next, we considered a first order Markov model in which the next bout in the sequence depends only on the last bout performed. Such a model outperformed the random model in predicting bouts following J-turns, orientations, “slow 1” swims, “slow 2” swims, and burst swims (**Figure 4B**; 30, 61, 29, 13, 106% improvement respectively). We subsequently built higher-order Markov models with a longer memory that considered multiple previous bouts in the sequence. Doing so continued to improve our ability to predict bouts following “slow 1” swims and capture strikes (14 and 4% improvement respectively), and “slow 2” swims, turns and burst swims (4, 12, 13% improvement respectively); but notably not those following J-turns and orientations (**Figure 4B**). From this analysis, we conclude that, during spontaneous swimming, previous bouts in a chain influence the future behavior of the animal. In contrast, during prey capture swimming, actions more than a single bout in the past have minimal observable influence on future bouts.

**Figure 4.**
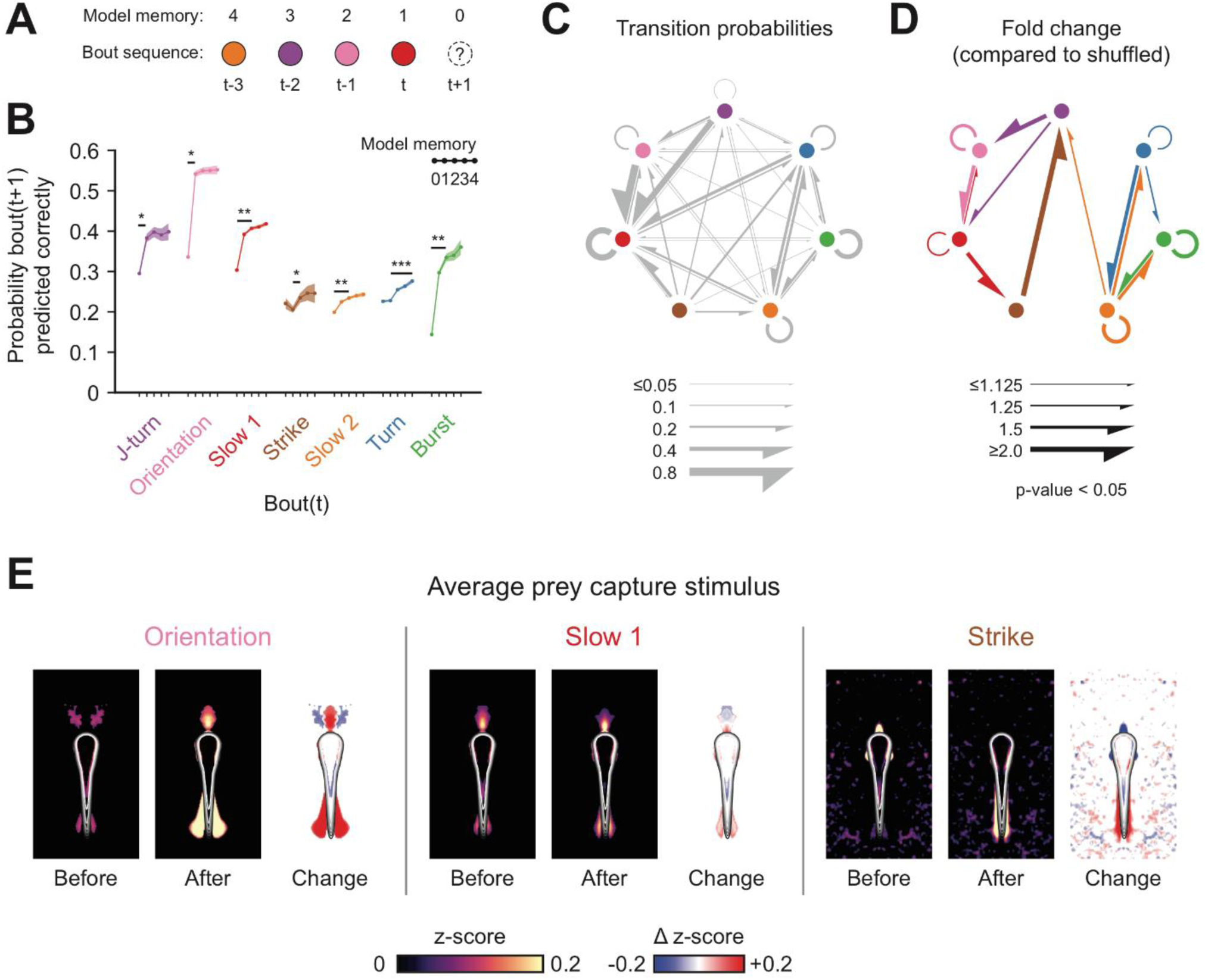
A stimulus-response loop drives predictable sequences through prey capture modules. (**A**) Example of a hypothetical bout sequence color-coded according to the behavioral module. We constructed a series of Markov models to predict the next bout in behavioral sequences. (**B**) Markov modelling of behavioral sequences. Five models were tested with different memory lengths of previous bouts in the sequence. Shaded areas represent 95% confidence interval across all occurrences of each module. Stars indicate improvement over previous model to predict future bouts (p value < 0.05, Student’s t-test, Holm-Bonferroni correction). (**C**) Ethogram of zebrafish behavior. Colored circles represent behavioral modules, gray arrows indicate transition probabilities between modules. (**D**) Transition probabilities significantly higher than chance (p-value < 0.05, Mann-Whitney U test, Holm-Bonferroni correction). Arrows show fold change in probability compared to shuffled data. (**E**) Average transformation of the visual scene produced by prey capture modules. Maps show average pixel intensity (prey) around fish across all bouts before (left) and after (middle) each module, expressed as a z-score (normalized using mean and standard deviation of 90,000 randomly selected frames). Difference is shown on the right. Images are thresholded using 95^th^ percentile. Contour shows outline of the fish. See also **Figure 4 – figure supplement 1**.

We next asked which specific behavioral transitions accounted for the stereotypy we observed in prey capture module chaining. We found that animals are more likely to transition from J-turns and orientations to “slow 1” forward swims than the reverse (**Figure 4C**). Transitions in the sequence, J-turn, orientation, “slow 1”, capture strike, were more than 1.5 times more likely to occur than expected by chance (**Figure 4D**). Moreover, we found the majority of transitions between prey capture and spontaneous modules were less likely than chance (18 / 24 transition pairs). Transitions within spontaneous modules (“slow 2” swims, burst swims and routine turns) were significantly overrepresented (6 / 6 transition pairs). However, in contrast to the stereotyped sequences we observed during prey capture, switching between spontaneous modules was more stochastic. We also found a high incidence of repetitive behaviors – performing the same module more than once successively in a behavioral chain (5 / 7 transitions to same module). Collectively, these results demonstrate a hierarchical organization to zebrafish behavior, with different modules and chaining dynamics underlying spontaneous and prey capture swimming.

### Prey capture sequences are maintained through tight stimulus-response loops

We reasoned that changes in the visual stimulus received by fish as they orients towards and approach prey might cause switching between behavioral modules during prey capture. If such changes are reproducible, they might form the basis of a stimulus-response chain, in which completion of one bout generates the appropriate stimulus for releasing the next bout. To test this, we reconstructed the visual experience of zebrafish performing prey capture sequences from our raw video data (see Methods). Doing so, we inferred the average stimulus that fish see before the onset of each behavioral module; and how the fish’s actions transform the visual scene (**Figure 4E**). Larvae initiate hunting sequences with a J-turn or orientation about 50% of the time (**Figure 3 – figure supplement 2**), and we find these bring the prey from the lateral to the anterior visual field. We found this new stimulus to be correlated with the onset of “slow 1” swims, which bring the prey to a stereotyped position in the near-anterior visual field. Prey in the near-anterior visual field was associated with the onset of capture strikes. Thus, the successive transformation of the visual scene as a result of the fish’s own motion could account for the stereotyped sequence through behavioral modules we observe during prey capture. In contrast, we do not observe stereotyped stimuli associated with spontaneous modules (**Figure 4 – figure supplement 1**), suggesting behavioral switching during this swimming state is likely mediated by internal neural processes.

### Prey capture chains conclude with a distance-dependent choice of strike type

Curiously, we noted that the most variable module in our data, the capture strike, seemed to be associated with the most stereotyped sensory stimulus – a paramecium in the near-anterior visual field. To investigate the source of this variation, we examined the prey capture strike further, with the goal of uncovering latent structure in this behavior masked by larger differences between bouts represented in our behavioral space. Our first hypothesis was that variation in capture strikes would be the result of a mixture of “long” and “short” capture dynamics (Marques et al., 2018). Capture strike durations clearly form a bimodal distribution, with one peak around 100 ms and a second peak around 200 ms (**Figure 5A**). Across all capture strike durations, however, we noticed that fish consumed the prey after a stereotyped time, and that long capture strikes resulted from a second, spontaneous-like bout being triggered immediately after the capture event. From this, we concluded that long capture dynamics were the result of bout concatenation, and so we hypothesized that variation in capture strikes was largely due to the post-capture phase.

**Figure 5.**
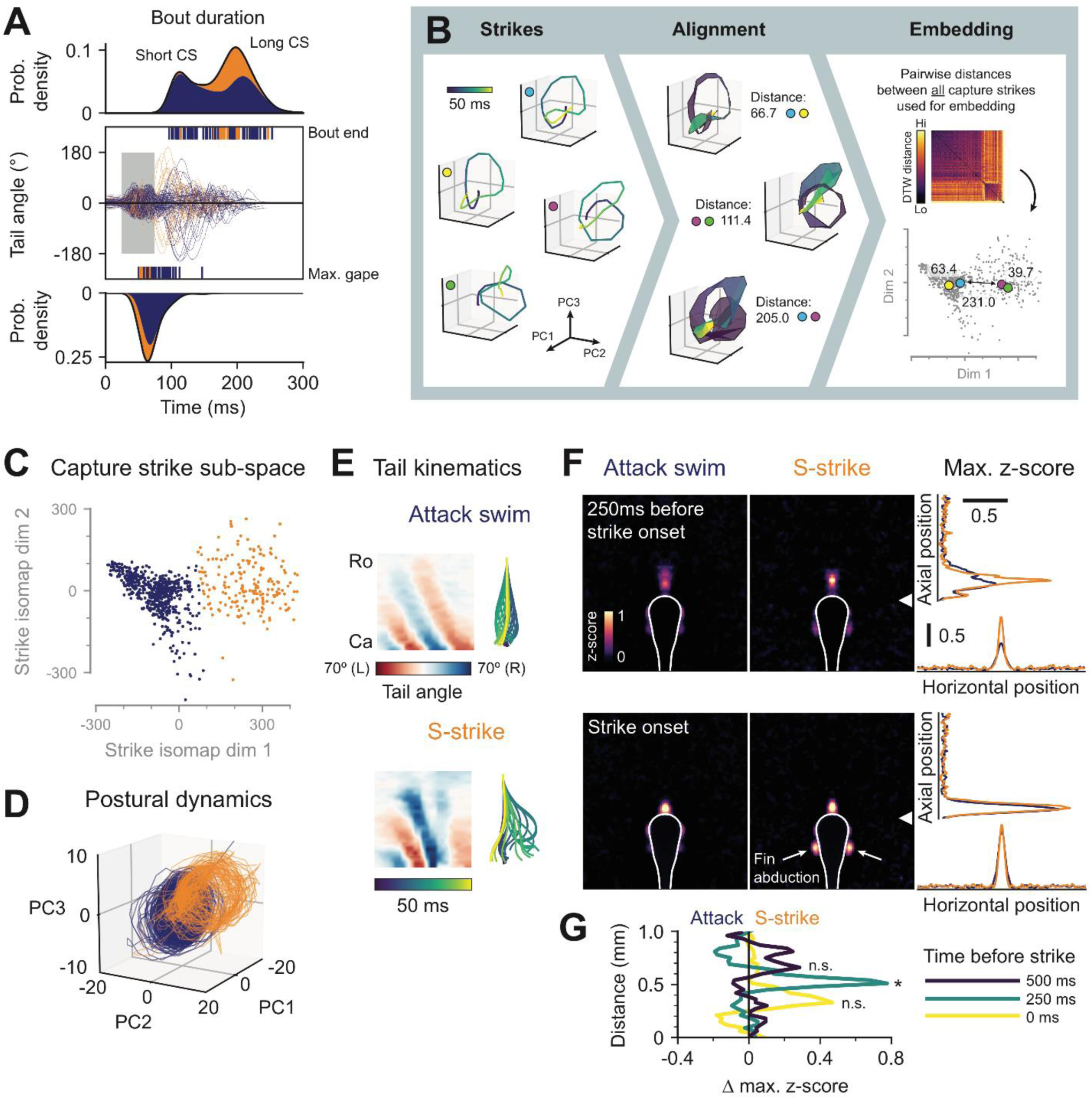
Zebrafish larvae perform distinct capture swims depending on prey position. (**A**) Captures strikes consist of a capture phase and a variable post-capture phase. Top: kernel density estimation of capture strike duration for attack swims (blue) and s-strikes (orange), shown as a stacked histogram. Middle: tail tip angle over time for capture strikes, defined by points in behavioral space containing > 50% late prey capture swims (see figure 2D). Grey window indicates initial 50 ms capture phase. Bottom: kernel density estimation showing when prey are consumed. (**B**) Pipeline for generating capture strike sub-space from initial capture phases. Strikes are represented by trajectories through postural space. Pairwise distances between all strikes, computed using DTW, are used to generate the sub-space with isomap embedding. (**C**) K-means clustering in the capture strike sub-space reveals two types of strike maneuver. (**D**) Trajectories through postural space for the two capture strike clusters. (**E**) Representative examples of an attack swim and an S-strike. Tail kinematics (left) and reconstructed bout (right). (**F**) Normalized average prey density around the fish 250 ms prior to (top) and at the onset of (bottom) each type of capture strike. White contour shows outline of fish. Bright spots either side of the fish contour at onset of S-strikes signify fin abduction (white arrows). Right: maximum prey density along axial and horizontal axes in the anterior visual field (measured from white arrowhead). (**G**) Difference in maximum prey density between S-strikes and attack swims as a function of distance from the fish 500 ms (dark blue), 250 ms (teal) and 0 ms (yellow) prior to strike onset. *p-value < 0.01, permutation test on the absolute maximum z-score difference; n.s. not significant.

To examine the stereotypy of the initial capture phase, we re-embedded capture strikes to produce a behavioral sub-space using our PCA-DTW-isomap pipeline, taking into account only a short 50 ms window before jaw opening (**Figure 5B**; see Methods). Doing so revealed two clearly separated clusters in the capture strike sub-space, suggesting that larvae capture prey with one of two distinct maneuvers (**Figure 5C**). These two clusters displayed markedly different postural dynamics (**Figure 5D**). We termed the two capture strike maneuvers the attack swim (blue cluster) and the S-strike (orange cluster) (**Figure 5E** and **Video 3**). S-strikes are immediately followed by a post-capture bout, possibly as a means to stabilize the animal in the water following the explosive capture maneuver. In contrast, only about half of attack swims lead into a post-capture bout (**Figure 5A**). These results reveal variation in bout dynamics exhibited by zebrafish larvae while striking at prey, suggesting that this behavior does not represent a single stereotyped movement, but rather two possible capture strategies employed by fish in different contexts.

To test if the two kinematically distinct capture maneuvers might be selected in response to different stimuli, we investigated the evolution of prey position around the fish over time for hunting sequences that resulted in either an attack swim or an S-strike, respectively (**Video 4**). We found prey position in the anterior visual field for the two types of strike started to diverge approximately 250 ms prior to the onset of the two maneuvers (**Figure 5F,G**). S-strikes occurred with a higher probability when prey was centered in the anterior visual field and 0.5 mm away within 250 ms of the onset of the swim (**Figure 5F,G**). For attack swims, prey became centered later in the bout chain and occurred within 0.25 mm of the fish. This difference was less prominent at the onset of the strike, suggesting that by this point the animal has already committed to one capture maneuver. In support of this, larvae characteristically abduct their pectoral fins prior to the onset of the S-strike but not the attack swim (McClenahan et al., 2012) (**Figure 5F**, white arrows). Together, these results indicate that the distance to the prey determines the choice of capture maneuver, with the S-strike recruited for prey located further than 0.25 mm, and the attack swim used to capture nearer prey.

### Variable tail kinematics combine with stereotyped jaw movements to capture prey from below

Fish must coordinate their tail movements during capture strikes with jaw movements that generate suction to draw the prey into their mouths (Hernández et al., 2002; Patterson et al., 2013). The degree of stereotypy in jaw movements is unknown, and it is possible that they, too, form discrete modules that are part of the prey capture chain. Therefore, we modified our recording setup and simultaneously observed tail and jaw kinematics of zebrafish larvae during prey capture (**Figure 6A,B**; see Methods). We tracked the position and pitch of larvae as well as the base of the jaw and elevation of the cranium (**Figure 6B,C** and **Video 5**). We found that the majority of jaw movements performed by zebrafish larvae were initiated immediately after a swim bout (**Figure 6D**), suggesting a stereotyped, sequential activation of these two types of movement. We then applied our PCA-DTW-isomap embedding pipeline to generate a behavioral space of jaw movements (**Figure 6E,F**). In this space, we could identify two well-separated clusters (**Figure 6F**). The larger cluster corresponds to a relatively slow, low amplitude depression of the lower jaw with little or no movement of the cranium (**Figure 6G**, left). Another type of jaw movement was rare but highly stereotyped, comprising a rapid, large amplitude depression of the lower jaw concurrent with cranial elevation (**Figure 6G**, right; **Video 6**). This movement was exclusively associated with attempts to capture prey. Inspecting the bouts preceding incidents of capture-associated jaw movements, we identified three distinct capture actions performed by zebrafish larvae (**Figure 6H**). These include S-strikes and attack swims, in addition to low-amplitude or absent tail movements corresponding to a purely “suction” capture (Hernández et al., 2002; Patterson et al., 2013). Thus, different capture strategies in zebrafish larvae emerge by combining variable tail kinematics with stereotyped jaw kinematics in a sequential chain.

**Figure 6.**
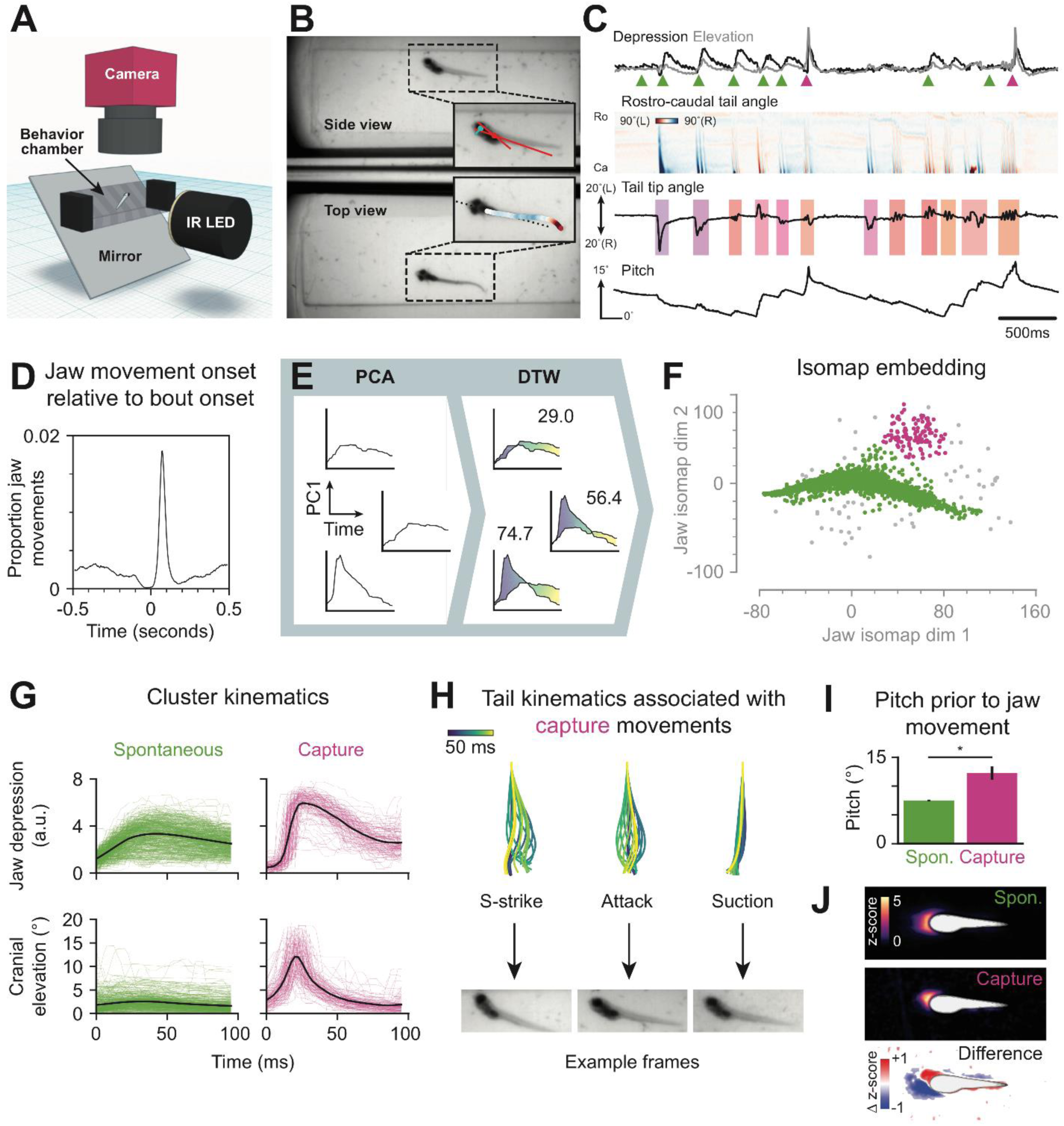
Capture strikes chain into a single stereotyped jaw movement. (**A**) Schematic of the setup used to record behavior simultaneously from above and from the side. (**B**) Example frame; insets are overlaid with tail and jaw tracking. (**C**) Jaw and tail kinematics from three seconds of behavioral recording. Top: depression of the jaw (black) and elevation of the cranium (gray). Arrowheads show onset of automatically identified jaw movements. Spontaneous movements (green); capture strikes (magenta). Middle: tail tracking. Bouts are color-coded according to nearest exemplar in behavioral space. Bottom: pitch of the fish in the water. (**D**) Cross-correlation between bout onsets and jaw movements onsets. (**E**) Pipeline for generating a jaw movement behavioral space. Jaw kinematics are transformed into postural dynamics with principal component analysis (PCA). Isomap embedding using DTW distances between jaw postural time series generates behavioral space. (**F**) HDSCAN clustering in the jaw behavioral space reveals two types of jaw movement. (**G**) Jaw depression (top) and cranial elevation (bottom) for the two types of jaw movement in zebrafish larvae. Colored traces: individual movements. Black lines: average. (**H**) Larvae can chain into capture jaw movements from three types of tail movement. Example of an S-strike, attack swim, and no tail movement that preceded the shown capture jaw movement (bottom). (**I**) Pitch of fish in the water prior to swims that chain into spontaneous and capture jaw movements. * two-tailed p-value < 0.01, unpaired Student’s t-test. (**J**) Normalized average prey density around the fish at the onset of bouts that chained into a spontaneous jaw movement (top) or a capture jaw movement (middle). White contour: outline of fish in average image. Anterior is left. Images are aligned and rotated so that the fish is in a horizontal position. Bottom: z-score difference between capture and spontaneous images. Red indicates higher density preceding a capture maneuver.

We observed that hunting episodes of zebrafish larvae were associated with both changes in pitch and moving up and down in the water column (**Figure 6C**). On average, larvae have a preferred orientation of 7° in the water and rotate to 12° prior to the onset of a capture, suggesting that fish adjust their pitch as well as their azimuth over the course of a hunting sequence (**Figure 6I**). Analyzing the prey position around the fish prior to the onset of captures revealed a preferred position in the immediate anterior and slightly dorsal visual field (**Figure 6J**). Such a configuration implies that capture strikes are initiated when prey fall on the temporal-ventral retina and that cranial elevation and jaw opening then create downward suction of prey into the up-turned mouth of the fish. Spontaneous jaw movements were associated with prey near the head of the fish, suggesting that these movements may serve olfactory or gustatory functions.

### Genetic disruptions of visual processing perturb behavioral chaining

Our results suggest that prey capture in zebrafish is maintained through a stimulus-response loop, which drives predictable transitions between behavioral modules. These transitions are triggered by changes in the visual stimulus the fish receives as the behavior progresses. Therefore, we hypothesized that genetic mutants with different visual impairments should have selective deficits in behavioral chaining during prey capture. In zebrafish larvae, prey capture depends on vision, and is impaired in darkness as well as in blind mutants (Gahtan et al., 2005; Patterson et al., 2013). Prey capture circuitry includes retinal ganglion cells (RGCs) and the optic tectum (Bianco and Engert, 2015; Gahtan et al., 2005; Semmelhack et al., 2014). In *lakritz* mutants (*lak*^*th241*^) (Kay et al., 2001; Neuhauss et al., 1999), RGCs fail to develop, and, consequently, these fish are blind (**Figure 7A**, bottom left). We reasoned that if vision drives transitions into and through prey capture, these swims should be absent in *lak-/-*. To test this, we recorded the behavior of *lak-/-* and sibling controls (mix of *lak+/-* and *lak+/+*) in the presence of prey and mapped their bouts into our canonical behavioral space (**Figure 7B**, top). We found a 58% reduction in the number of prey capture bouts performed by mutants compared to controls (**Figure 7C**, left; controls, 39.3% ± 0.04, n=6; mutants, 16.5% ± 0.04, n=6; mean ± SD). This could be explained by a decreased probability of initiating prey capture modules in mutants, as well as a failure to sustain sequences for more than a single bout once initiated (**Figure 7D,E**; spontaneous sequence lengths: controls 1.72 ± 0.08, mutants 2.23 ± 0.18; prey capture sequence lengths: controls 2.06 ± 0.13, mutants 1.38 ± 0.10; mean ± SD). These differences were reflected in the SVD of the transition frequency matrices of controls and mutants (**Figure 7 – figure supplement 1A-E**). While the first two symmetric and first antisymmetric transition modes of controls closely matched wildtypes in the canonical dataset (**Figure 7 – figure supplement 1B,C**, dot products 0.96, 0.88 and 0.72, respectively), transition modes involving prey capture swims were disrupted in mutants (**Figure 7 – figure supplement 1D,E**, dot products 0.59, 0.13 and 0.06). Thus, depriving animals of visual inputs selectively disrupts the initiation of prey capture modules.

**Figure 7.**
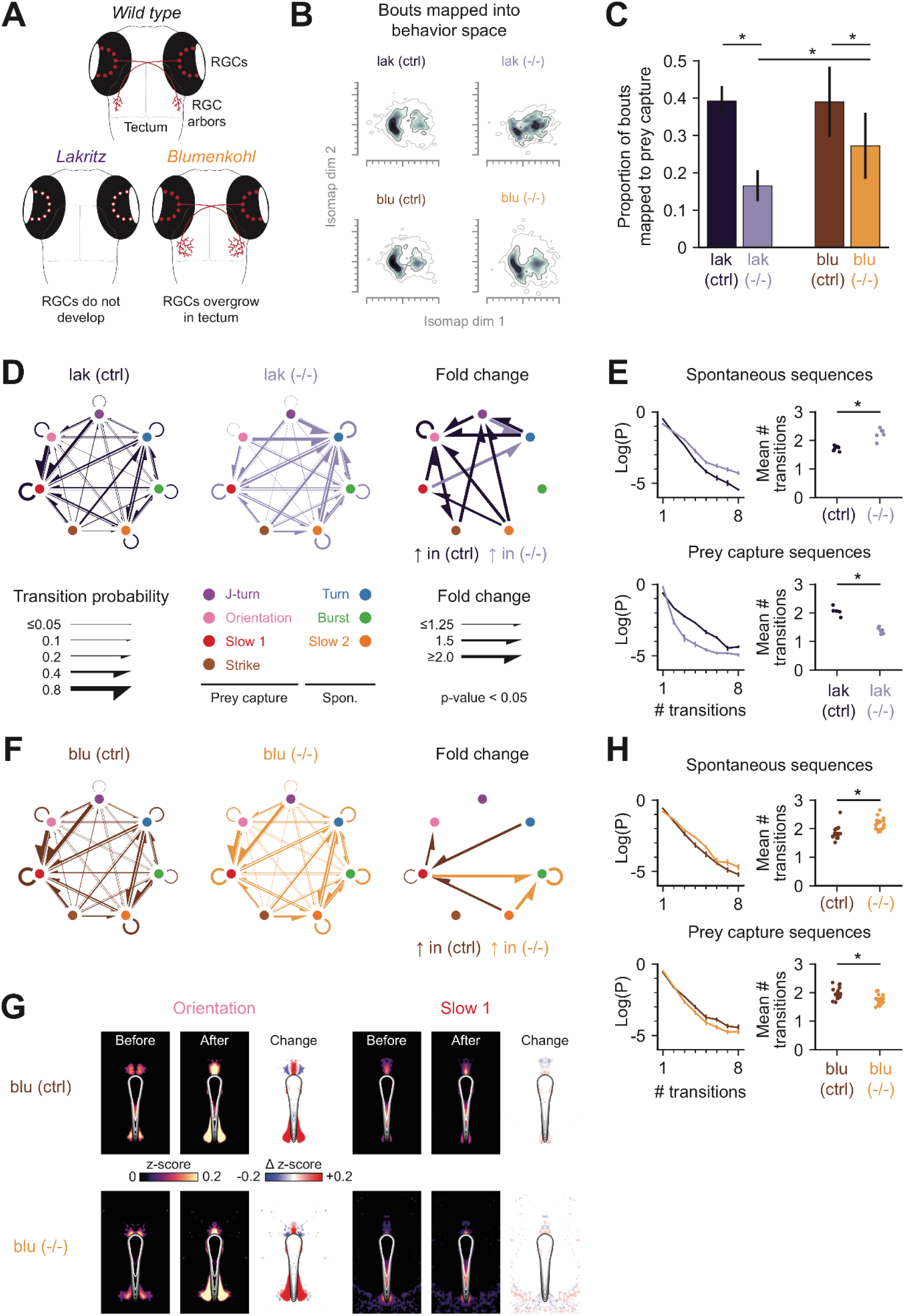
Behavioral chaining is disrupted in *lakritz* and *blumenkohl* mutants. (**A**) Developmental phenotype of *lakritz* (*lak*) and *blumenkohl* (*blu*) mutants. In *lak-/-* RGCs fail to form. In *blu-/-* RGC arbors overgrow in the tectum. *Lak* controls are a mix of *lak+/-* and *lak+/+* siblings. *Blu* controls are *blu+/-* siblings. (**B**) Bouts from *lak* and *blu* can be mapped into the canonical behavioral space using DTW, enabling comparison of different behavioral datasets. (**C**) Proportion of bouts that are mapped to prey capture exemplars. Error bars indicate standard deviation. * two-sided p value < 0.01 Mann-Whitney U test. Difference between control groups is not significant (p > 0.4). (**D**) Ethogram of *lak* behavior. Controls (left), mutants (middle), and fold change in transition probabilities between these groups (right). Only statistically significantly differences between groups are shown (p < 0.05 Mann-Whitney U test with Holm-Bonferroni correction). (**E**) Spontaneous (top) and prey capture (bottom) sequences in *lak*. Left: probability that sequence is aborted after a given number of bouts. Right: average sequence length. * two-sided p value < 0.01 Mann-Whitney U test. (**F**) Ethogram of *blu* behavior. Controls (left), mutants (middle), and fold change in transition probabilities between these groups (right). Only statistically significantly differences between groups are shown (p < 0.05 Mann-Whitney U test with Holm-Bonferroni correction). (**G**) Normalized average prey density before and after orientations and “slow 1” swims for *blu* controls (top) and mutants (bottom). Images are thresholded using 95^th^ percentile. (**H**) Spontaneous (top) and prey capture (bottom) sequences in *blu*. Left: probability that sequence is aborted after a given number of bouts. Right: average sequence length. * two-sided p-value < 0.01 Mann-Whitney U test. See also **Figure 7 – figure supplement 1**.

Next, we tested the behavior of *blumenkohl* mutants (*blu*^*tc257*^) (Neuhauss et al., 1999), which carry a mutation in a vesicular glutamate transporter, *vglut2a. Blu-/-* mutants grow larger RGC axonal arbors in the tectum, which is proposed to decrease visual acuity in these animals (**Figure 7A**, bottom right) (Smear et al., 2007). Consequently, *blu-/-* mutants are less efficient hunters of small prey items. According to our model, these mutants should be able to initiate prey capture sequences, but we predicted their blurred vision would prevent them from receiving the appropriate stimuli required to connect subsequent bouts in the behavioral chain. We found that both *blu-/-* and *blu+/-* sibling controls exhibited the full behavioral repertoire of wild-types (**Figure 7B**, bottom); however, mutants performed 30% fewer prey capture bouts compared to controls (**Figure 7C**, right; controls, 39.0% ± 0.09, n=18; mutants, 27.2% ± 0.09, n=19; mean ± SD). The transition modes of controls were indistinguishable from wildtype (**Figure 7 – figure supplement 1F-H**, dot products 0.95, 0.93 and 0.87), but were disrupted in mutants, suggesting that behavioral chaining was affected in these animals (**Figure 7 – figure supplement 1F,I,J**, dot products 0.64, 0.08, 0.10).

We found the most significant changes in the *blu-/-* behavior affected transitions into and out of “slow 1” swims, recruited during prey capture, and burst swims, recruited during spontaneous swimming (**Figure 7F**). We predicted that blurred vision in the mutants could prevent them from receiving the appropriate stimulus necessary to initiate “slow 1” swims during prey capture. Therefore, we investigated the prey position around *blu-/-* animals during hunting sequences. This revealed that *blu-/-* mutants perform orientations when prey were closer to the animals than in controls (**Figure 7G**, left). We also found that prey were closer prior to the onset of “slow 1” swims in *blu-/-*, and was in a less reproducible location (**Figure 7G**, right). We reasoned that the nearer position required to orient towards prey in mutants would result in them initiating prey capture less frequently, and indeed we found that *blu-/-* perform more spontaneous bouts before initiating a prey capture swim (**Figure 7H**, top, spontaneous sequence lengths: controls 1.84 ± 0.22, mutants 2.16 ± 0.2; mean ± SD). Second, we predicted that the less stereotyped prey position prior to “slow 1” swims would impair mutants’ ability to maintain prey capture sequences. Indeed, we found that prey capture sequences in *blu-/-* were slightly truncated (**Figure 7H**, bottom, prey capture sequence lengths: controls 1.97 ± 0.20, mutants 1.75 ± 0.16; mean ± SD). Thus, our fine-grained analysis reveals a specific deficit in visually driven chaining of prey capture sequences that likely results from a blurred visual map in the optic tectum.

## Discussion

A thorough quantification of the actions performed by an animal, and how the environment influences these actions, is a prerequisite for understanding the sensorimotor transformations performed by the brain to realize behavior (Krakauer et al., 2017). Our unsupervised analysis reveals that zebrafish swim bouts lie on a behavioral continuum. We group bouts into seven behavioral modules that are differentially recruited during spontaneous swimming and prey capture. During prey capture, we find that the location of prey in the visual scene likely triggers a specific behavioral module, whose movement transforms the prey stimulus, leading to the next bout in the chain. Thus, iterative bout chaining positions the prey in the center of the anterior dorsal visual field through a stimulus-response loop. Once the prey has reached this position, the fish releases one of three distinct capture maneuvers, determined by the remaining distance to the prey. Further decomposition of these capture maneuvers revealed that distinct predation strategies arise through combining a stereotyped jaw movement with distinct types of swim bout. Genetic manipulation of visual processing disrupted the stimulus-response loop, preventing prey capture initiation in blind animals, and impairing the maintenance of the behavior in animals with blurry vision.

One of the challenges of linking behavior to neural activity is finding suitable representations that link these two domains (Brown and Bivort, 2018). Postural modes identified through PCA have previously been used to describe spontaneous swimming in zebrafish and crawling behavior in *C. elegans* and *Drosophila* larvae (Girdhar et al., 2015; Stephens et al., 2008; Szigeti et al., 2015). We found that the first three eigenfish in our data form a harmonic series (**Figure 1E**), with the second and third modes describing the sinusoidal oscillation of the tail during a bout and the first mode accounting for turning. These are similar to the basis vectors used to describe postural dynamics in fly maggots and nematodes, suggesting that such modes may serve as a common framework for finding equations of motion across taxa. In the future it may be possible to relate trajectories in this space to specific structural or dynamical motifs in the neural circuits that produce locomotion across species (Kato et al., 2015).

Different unsupervised approaches have not provided a consensus on whether behavior is organized into distinct modules with stereotyped kinematics (Berman et al., 2014; Marques et al., 2018) or whether such modules represent extremes of a continuum (Katsov et al., 2017; Szigeti et al., 2015). Non-linear embedding algorithms, such as t-SNE, have become popular for analyzing high-dimensional behavioral data (Berman et al., 2014; Marques et al., 2018); however, such representations separate behaviors along arbitrary dimensions and tend to exaggerate the distances between clusters. Our isomap embedding approach revealed a continuum of swim bouts used by zebrafish larvae during hunting and spontaneous swimming (**Figure 2B**). This showed that swim bouts predominantly vary in swimming speed and turning degree (**Figure 2 – figure supplement 1C**). When we inspected the prey capture strike further, however, we found evidence of modularity in this particular behavior (**Figure 5**). Moreover, we used our method to explore the kinematics of jaw movements, demonstrating that it generalizes to different types of movement patterns (**Figure 6**). In addition, we demonstrate that different datasets can be bridged into the same behavioral space, aiding the identification of behavioral deficits in mutants (**Figure 7**). Thus, our method provides a suitable alternative to stochastic embedding algorithms as it can capture both continuity and discreteness in a variety of behavioral datasets.

Classically, two general models have been proposed to explain how animals chain behavioral modules into sequences. Hierarchical models propose that behavioral switching is stochastic, yet structured over various timescales (Berman et al., 2016; Seeds et al., 2014). In contrast, sequential models predict recurring, stereotyped behavioral chains (Long et al., 2010). These are not mutually exclusive, and it has been suggested that both might contribute to spontaneous behaviors in *Drosophila* (Berman et al., 2016; Katsov et al., 2017). The extent to which these mechanisms contribute to the production of behavioral sequences in zebrafish was not known. We found that larvae preferentially form sequences consisting of either spontaneous or prey capture modules (**Figure 3C,D** and **Figure 4C,D**), suggesting a hierarchical organization in their behavior. We found bout chains to be more stereotyped during prey capture, hinting that an underlying mechanism drives sequential activation of modules during this behavior (**Figure 2D, Figure 3D** and **Figure 4B-D**). Predictable behavioral sequences can be driven by either internal neural mechanisms or feedback from the environment (Coen et al., 2014; Long et al., 2010). Previous reports have suggested that zebrafish larvae reflexively react to the current position of a prey object in the visual field when generating their bouts (Patterson et al., 2013; Trivedi and Bollmann, 2013). Our results suggest that a stimulus-response loop links successive bouts in a behavioral chain and drives stereotyped sequences through prey capture modules, with little influence from previous behaviors (**Figure 4**). Short integration windows for deciding the next behavioral action have also been observed in thermal navigation of larvae (Haesemeyer et al., 2015) and social affiliation of juvenile zebrafish (Larsch and Baier, 2018). Thus, stimulus-response loops driving behavioral chaining might not be specific to prey capture, but provide a more general mechanism underlying goal-directed behavior in zebrafish.

Zebrafish larvae move in a three dimensional water column, and make full use of this environment during natural behaviors (Horstick et al., 2017). It was recently proposed that a specialized UV-sensitive zone in the ventral retina could facilitate targeting prey from below (Zimmermann et al., 2018). We demonstrate that larvae do indeed orient themselves beneath the prey over the course of a hunting sequence (**Figure 6I,J**). Furthermore, we found that larvae capture prey with a single stereotyped jaw movement that includes dorsal flexion of the cranium (**Figure 6F-H, Video 5** and **Video 6**). This movement likely generates downward suction during strikes. Moreover, this jaw movement is either produced in isolation, or in combination with an attack swim or S-strike maneuver, both of which are similarly stereotyped and occur when prey reach a specific location in the visual field (**Figure 5C-G** and **Video 4**). These results provide compelling evidence that jaw morphology has co-evolved with sensory and motor circuits to reduce the complexity of capturing prey in a three-dimensional environment. Producing invariant actions in response to stereotyped “releasing” stimuli has long been considered an efficient way to ensure reproducible outcomes in innate behaviors (Ewert, 1987; Tinbergen, 1951). By linking three different releasing stimuli to three stereotyped motor programs, all sharing a common jaw movement, the developing nervous system of the zebrafish larva has evolved an efficient means to produce reliable, flexible behavior with a limited number of neurons.

Whether behaviors exist in a continuum or as stereotyped, invariant motor patterns have different implications for their underlying neural circuit implementation. Behavioral continua, such as the J-turns, orientations and “slow 1” swims that occur during prey capture, may be encoded in a topographic motor map, where the position of prey in the visual field is transformed into a graded motor output. Such a map has been identified in the optic tectum of zebrafish larvae and its projections to the hindbrain (Helmbrecht et al., 2018). We found that *blu* mutants, which have blurred retinotectal maps, had difficulty sustaining prey capture sequences. This suggests that the visuo-motor transformations normally performed by the tectum during prey capture are disrupted in these mutants. Furthermore, the gradual transformation of the visual stimulus received by fish over a prey capture sequence suggests that the animal is trying to position the prey at a specific point in the temporal-ventral retina (**Figure 4E, Figure 5F,G** and **Video 4**). When the eyes are converged, this point would result in the prey being represented bilaterally in the anterior regions of both tecta. This region could contain specialized circuitry for implementing the appropriate capture maneuver, depending on the distance to the prey. Rather than a continuous motor map, we posit the S-strike and attack swim are driven by separate command-like neuronal populations, or alternatively by different activity levels within a common population. Similarly, a dedicated neural circuit may control the stereotyped jaw movement we observe during strikes (**Figure 6** and **Video 6**). Thus, we propose that different neural architectures underlie the pursuit and capture of prey in zebrafish larvae. Our work provides a computational framework for interrogating the production and chaining of motor modules during this behavior in a genetically tractable vertebrate.

## Methods

### Fish

For experiments relating to **Figure 1-6** we obtained TLN (nacre) embryos from an outcross of TLN homozygous to TL/TLN heterozygous adults. Until 3 days post fertilization (dpf) embryos were raised in Danieau’s solution (17 mM NaCl, 2 mM KCl, 0.12 mM MgSO_2_, 1.8 mM Ca(NO_3_)_2_, 1.5 mM HEPES) at a density of 60 embryos per 50 ml at 28 °C with a 14h-10h light-dark cycle. Thereafter, embryos were transferred to new dishes containing fish system water and raised at a density of 30 larvae per 50 ml until behavioral testing at 7 dpf or 8 dpf. At 5 dpf and 6 dpf, a few drops of dense paramecia culture (*Paramecium multimicronucleatum*, Carolina Biological Supply Company, Burlington, NC) were added to each dish and larvae were allowed to feed *ad libitum*.

For experiments relating to **Figure 7**, we used *lakritz* (*lak*^*th241*^) and *blumenkohl* (*blu*^*tz257*^) mutants (Neuhauss et al., 1999) in a TL background. *Lak* mutants were obtained from a heterozygous in cross. Homozygous mutants could be clearly identified by their dark color compared to sibling controls (mixture of heterozygotes and wild types) in a visual background adaptation (VBA) assay. *Blu* mutants were obtained by outcrossing heterozygous females to homozygous males. Similarly to *lak*, mutants could be identified unambiguously with a VBA assay. Larvae were raised as described above, except they were not fed at 5 and 6 dpf, and thus their naïve prey capture ability was assayed at 7 dpf. This was to minimize potential confounding effects of experience-dependent improvement in prey capture efficacy between groups.

### Free-swimming behavioral assay

Free-swimming prey capture experiments relating to **Figure 1-5** and **Figure 7** were conducted using a custom-built behavioral setup. Behavior arenas were produced by flooding a 35 mm petri dish with 2% agarose (Biozym, Germany), with an acrylic square (15 × 15 mm, 5 mm deep) placed in the center. Once the agarose had set, the acrylic square was removed producing a hollow chamber with transparent walls. Single larvae were introduced to the chamber along with a drop of culture containing approximately 50-100 paramecia. The chamber was filled to the top with fish system water and a glass coverslip was placed over the chamber to flatten the meniscus. This provided a clean, transparent chamber where behavior could be observed and tracked.

Behavior experiments were performed in a climate-controlled box kept at 28 ± 1 °C between 3 and 12 hours after lights on. Each larva was recorded for 20 minutes using a high speed camera (PhotonFocus, MV1-D1312-160-CL, Switzerland), fitted with an objective (Sigma 50 mm f/2.8 ex DG Macro, Japan), connected to a frame grabber (Teledyne DALSA X64-CL Express, Ontario, Canada). The camera was positioned over the behavior arena, which was lit from below with a custom-built infrared LED array. Behavior was filmed at 500 frames per second with a frame size of 500 × 500 pixels covering an area slightly larger than the arena (**Figure 1B**), providing a final resolution of approximately 0.03 mm/pixel. The aperture of the camera objective was adjusted such that the fish was in focus throughout the entire depth of the arena. Recording was performed using StreamPix 5 software (NorPix, Quebec, Canada) and individual trials were initiated through a custom written Python script. Each 20 minute session was split into 20× 1 minute recording trials, with < 1 second between the end of one trial and the beginning of the next, to keep video files to a manageable size. If frames were dropped during a trial, the recording was stopped to prevent problems in subsequent analyses. Videos were compressed offline in VirtualDub with Xvid compression before tracking was performed.

### Free-swimming behavioral assay in three dimensions

To record behavior simultaneously from above and from the side, we designed a new chamber (**Figure 6A**). A 3 ml transparent, unfrosted plastic cuvette was with flooded with 2% agarose. An acrylic rod (20 × 5 × 5 mm) was inserted into the liquid agarose, which was allowed to set, after which the rod was removed leaving behind a hollow chamber. As before, individual larvae were introduced into the chamber with a drop of paramecia culture topped up with fish system water. The opening was plugged with a small piece of acrylic cut to match the cross section of the chamber (5 × 5 mm). The cuvette was placed on its side on top of a glass coverslip suspended above a mirror angled at 45°. The high speed camera was positioned above this setup in such a way as to allow the fish in the chamber as seen from above as well as the reflected side view from the mirror to be visible within the field of view of the camera. The IR LED array was rotated by 90°, allowing the chamber to be illuminated from the side and from below (via the mirror) with a single light source. We reduced the aperture of the camera objective so that the entire arena was in focus in both views and offset the decrease in luminance by increasing the exposure time of each frame. Consequently, for this experiment we achieved a frame rate of 400 fps. As described above, data from each fish was split into 20× 1 minute recording trials.

To record jaw movements during prey capture with higher spatial resolution in **Video 6**, we used two cameras (PhotonFocus, MV1-D1312-160-CL, Switzerland) and two light sources and filmed a number of fish swimming in a custom-built transparent chamber. We waited for one of the fish to start hunting a paramecium in the field of view of both cameras and manually triggered the recording. Frame acquisition was synchronized using StreamPix 5 and a dual camera frame grabber.

### Tail and eye tracking

Tracking was performed using custom-written Python scripts. Each frame was tracked independently. Each frame was divided by a background image, calculated as the median of every 100^th^ frame over all trials from a given animal. The frames were then thresholded and contours extracted using OpenCV. The largest contour in the image was taken as the outline of the fish and all other pixels were discarded. Then, the histogram of pixel values of the fish was normalized and a second threshold was applied to find the three largest contours within the fish, corresponding to the two eyes and swim bladder. The eyes were identified automatically as the two contours with the nearest centroids and left and right identities were assigned using the sign of the vector product between lines connecting the swim bladder to these two points. The heading of the fish was defined by a vector starting in the center of the swim bladder and passing through the midpoint between the eye centroids. The angle of each eye was calculated from the image moments of their contours and was defined as:

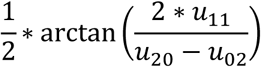

Where *u*_*ij*_ is the corresponding central moment. The eye angles in an egocentric reference were calculated as the difference between the heading angle and absolute orientation of eyes, and eye convergence defined as the difference between the eye angles (**Figure 2 – figure supplement 2A**). A 100 ms median filter was applied to smooth the traces obtained from each eye while preserving edges. The two thresholds used for tracking were set manually for each fish. In frames where the eye contours could not be detected through thresholding, we instead applied a watershed algorithm to obtain contours and then proceeded as above.

Due to the dark pigmentation of *lak* and *blu* mutants, there was insufficient contrast to segment the eyes from the surrounding skin using either thresholding or watershed analysis. For this reason, eye tracking could not be performed in these animals. To calculate the heading in this case, we used the second threshold to segment the head and body of the fish from the tail, for which we identified the minimum enclosing triangle using OpenCV. The heading was then defined as a vector passing through the apex and centroid of this triangle, and the position of the swim bladder estimated as lying midway between these two points.

To track the tail of the fish, we skeletonized the contour obtained after applying the first threshold described above. We started the tracking from the point on this skeleton nearest to the swim bladder. We used a custom-written algorithm to identify the longest path through the skeletonized image that started at this point, ended at the tip of a branch, and began in the opposite direction of the heading vector. We then linearly interpolated 51 equally spaced points along this path to obtain the final tail points.

The tail tip angle was defined as the angle between the midline of the fish (provided by the heading vector) and a vector between the center of the swim bladder and the last point of the tail. This angle is used to help visualize the sinusoidal oscillation of the tail in **Figure 1, 3, 5**, and **6**, but was not used as the basis of any analysis in the paper.

We vectorized the tracked tail points for kinematic analysis in a similar manner to what has been previously described (Girdhar et al., 2015; Stephens et al., 2008). Briefly, we calculated the angle between the midline (defined by the heading vector) and a vector drawn between each adjacent pair of tail points, providing a 50 dimensional representation of the tail in each frame. A three frame median filter was applied to the heading angle and tail kinematics to remove single frame noise.

The mean tail tip curvature was computed as the mean of the last ten points of the tail angle vector, and was used for bout segmentation. Bouts were detected by applying a threshold to the smoothed absolute value of the first derivative of this mean tail tip curvature. Uncharacteristically long bouts detected with this method were further split by finding turning points in the smoothed absolute value of the mean tail tip curvature convolved with a cosine kernel.

### Jaw tracking

As for the single view setup, each frame was tracked independently offline using custom-written Python scripts. Each frame was divided by a background image, calculated as the median of every 100^th^ frame over a recording trial. The upper and lower halves of the frame were tracked separately. The lower half of the frame, containing the image of the fish as seen from above, was tracked as described above. Fish were only tracked from the side when their heading was within ±45° of the imaging plane to minimize artifacts arising as a result of foreshortening. Frames were thresholded and contours extracted using OpenCV. The largest contour in the image was taken as the outline of the fish and all other pixels were discarded. Then, the histogram of pixel values of the fish was normalized and a second threshold was applied to find a contour enclosing the head and body of the fish. The pitch and angle of the cranium were calculated using image moments of these two contours respectively, with cranial elevation defined as the difference between them.

To find the point of the base of the jaw, we first defined point, ***p***, as the centroid of the head-body contour and vector, ***v***, defined by the cranium angle (i.e. orientation of this contour in the frame). We extended vector ***v*** from ***p*** until it intersected the fish contour at point ***q***. Next, we found the midpoint of 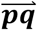, called ***c***. We then extended a vector orthogonal to ***v*** from ***c*** until it intersected the fish contour at the base of the jaw, ***h***. Jaw depression was defined as the Euclidean distance, 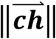.

The cranial elevation angle and hyoid depression were smoothed with an edge-preserving five-frame median filter. Then, we applied a high-pass filter by subtracting the baseline of these two kinematic features over a recording. To compute this baseline, we first calculated a 250 ms rolling minimum, and then computed the one-second rolling mean of this rolling minimum. This provided a relatively stable baseline for identifying jaw movements, despite changes in elevation and azimuth of the fish over a recording. To segment jaw movements, we identified periods when the baseline-adjusted jaw depression, smoothed with a 50 ms rolling average, was above a predetermined threshold and defined movement onset and offset as turning points in this smoothed trace.

### Embedding postural dynamics in a behavioral space

To generate our behavioral space, we excluded any bouts during which the tail of the fish hit the wall of the behavior chamber. This was to ensure that only the fish’s self-generated motion – and not motion artifacts introduced from distortion of the tail by the wall – was considered when mapping the behavioral space. Consequently, not all the bouts we observed could be mapped into the space.

To describe bouts in terms of their postural dynamics, we performed principal component analysis (PCA) on the tail kinematics across all bout frames. Data were normalized before applying PCA by subtracting the mean tail shape and dividing by the standard deviation.

The next step in generating the behavioral space involved computing the distance between every pair of bouts with dynamic time warping (DTW) (Sakoe and Chiba, 1978). DTW finds an alignment between two time series that minimizes a cost function, which is the sum of the Euclidean distances between each pair of aligned points. In our analysis, we only allowed trajectories to be warped within a 10 ms time window. For bouts of different lengths, we padded the end of the shorter bout with zeros until it was the same length as the longer bout. We performed each alignment twice, reversing the sign of all the values for one of the trajectories the second time, and considered the distance between two bouts to be: *min*(*DTW*(*t*_1_, *t*_2_), *DTW*(*t*_1_, −*t*_2_)), thus effectively ignoring the left/right polarity of the bouts.

For generating the behavioral space in **Figure 2**, we performed a round of affinity propagation (Frey and Dueck, 2007) prior to embedding, using the negative DTW distance between a given pair of bouts as a measure of their similarity. We used the median similarity between bouts as the preference for the clustering. Doing so provided 2,802 clusters, of which we excluded any clusters containing fewer than three bouts, thus ensuring that only repeatedly observable motor patterns were used for generating the behavioral space. As a final quality check, we manually inspected every cluster exemplar and removed incorrectly identified bouts, which usually was the result of tracking artifacts from a paramecium crossing the tail of the fish. The final number of clusters that we embedded was 1,744.

Since affinity propagation identifies an exemplar to represent each cluster, we produced our final behavioral space by performing isomap embedding (Tenenbaum et al., 2000) of these exemplars. For the isomap embedding, we constructed a nearest-neighbors graph of the exemplars using their DTW distances, and calculated the minimum distance between each pair of points in this graph. The isomap components correspond to the eigenvectors of this graph distance matrix.

### Eye convergence analysis

To identify periods of eye convergence, we calculated a kernel density estimation (Gaussian kernel, bandwidth=2.0) of the eye convergence angles across all frames for a given fish. This distribution was bimodal (eyes converged or unconverged) and therefore we defined the eye convergence threshold as the antimode (least frequent value between the two modes). To identify spontaneous, early, mid, and late prey capture bouts, we calculated the mean eye convergence angle over the first and last 20 ms of a bout, and concluded the eyes were converged if this number was above the threshold. Bouts were classified as spontaneous if the eyes were unconverged at the beginning and end of a bout; early prey capture if the eyes were unconverged at the beginning and converged at the end of the bout; mid prey capture if the eyes were converged at the beginning and end of the bout; and late prey capture if the eyes were converged at the beginning and unconverged at the end of the bout.

### Mapping kinematic features and eye convergence into the behavioral space

With our PCA-DTW-isomap approach, each point in the behavioral space represents a small cluster of bouts. For each bout, we calculated the mean speed, angle through which the fish turned, maximum angular velocity of the fish, and the time at which the maximum angular velocity occurred (turn onset). In **Figure 2 – figure supplement 1C**, we show the median of each of these features over a cluster. Similarly, we could calculate the proportion of bouts in each cluster that occurred during spontaneous, early, mid, or late prey capture as defined above. The prey capture index was defined as:

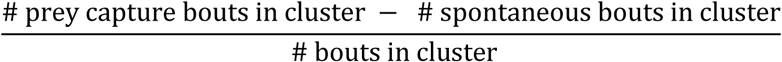

### Mapping mutant bouts into the behavioral space

To map mutant bouts into the behavioral space, we extracted tail kinematics and identified bouts as described above (see *tail and eye tracking*). The postural dynamics of each mutant bout was projected onto the first three principal components obtained from the canonical dataset (**Figure 1D,E**) to bring it into the same space as bouts from that dataset. Then, each mutant bout was mapped to one of the 1,744 exemplars identified in “*embedding postural dynamics in a behavioral space”* using dynamic time warping (DTW), with the nearest exemplar having the smallest DTW distance to the bout. In this way, each mutant bout could be projected into the three dimensional behavioral space defined by the 1,744 exemplars. In **Figure 7B**, we show a kernel density estimation of all bouts from a given condition over the first two dimensions of the behavioral space.

Since we could not perform eye tracking in the mutants (see *tail and eye tracking*), we instead calculated the proportion of bouts performed by each fish that were mapped to a prey capture motif, defined as those having a prey capture index > 0. This provided each fish with a “prey capture score”. We then compared the prey capture scores of fish with different genotypes with three two-sided Student’s t-tests (independent samples) comparing *lak* controls to *lak* mutants, *blu* controls to *blu* mutants, and *lak* mutants to *blu* mutants.

### Singular value decomposition (SVD) of behavioral transitions

To identify transition modes, we generated a transition frequency matrix, *M*, where *M*_*ij*_ contains the number of transitions from behavioral motif *j* to behavioral motif *i*, where each behavioral motif is a small cluster of bouts in the behavioral space (see *embedding postural dynamics in a behavioral space*). This matrix included all the transitions from all animals for a given experiment.

Since there are more than 3 million (1,744^2^) possible transitions between motifs, and only 44,154 transitions in our largest dataset, the matrix *M* is necessarily sparse. This would hinder the identification of common dynamical motifs, and so we performed smoothing on matrix *M* by blurring similar transitions into each other. To achieve this, we took advantage of the fact that nearby points in our behavioral space encode bouts with similar postural dynamics. We computed a weighting matrix, *W*, where 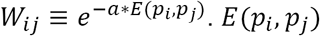 is the Euclidean distance between a pair of points in the three-dimensional behavioral space, and *a* is a smoothing factor (see **Figure 3 – figure supplement 1**).

We normalized matrix *W* so that the columns summed to one and then smoothed the transitions in matrix *M* with the transformation: *M*_*smooth*_ = *WMW*^*T*^.

To distinguish between symmetric transitions (i.e. those that occur in both direction), and antisymmetric transitions (i.e. those in which transitions in one direction outweigh those in the other), we decomposed the smoothed matrix, *M*_*smooth*_, into its symmetric and antisymmetric parts, where:

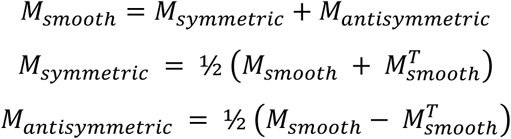

The symmetric and antisymmetric transition modes were found by taking the SVD of these two matrices respectively.

Every real or complex matrix, *A*, can be factorized using the singular-value decomposition (SVD) into three matrices such that:

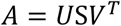

The columns of *U* and rows of *V*^*T*^ define two sets of orthonormal basis vectors and S is a diagonal matrix containing the singular values, ordered from largest to smallest. The SVD describes the transformation performed by matrix, *A*. Under this transformation, each row of the matrix, *V*^*T*^, is mapped to the corresponding column of *U* and scaled by the associated singular value. Therefore, this decomposition provides an unbiased description of the most common transitions between behavioral motifs.

A symmetric matrix, such as *M*_*symmetric*_, geometrically defines a scaling transformation. Consequently, its singular-value decomposition is the same as its eigendecomposition: spaces *U* and *V* are the same and S contains the eigenvalues. As such, the *n*^*th*^ transition mode of *M*_*symmetric*_ can be written:

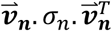

Where 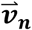 is the singular vector with corresponding singular value, *σ*_*n*_. In **Figure 3C** and **Figure 7 – figure supplement 1**, we show motifs with positive or negative loadings to each singular vector separately.

An antisymmetric matrix, such as *M*_*antisymmetric*_, describes a set of orthogonal rotations. As such, spaces *U* and *V* are related by a 90° rotation and each transition mode can be written:

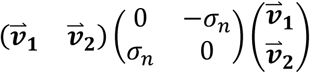

Where 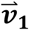 and 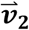 are orthonormal, and *σ*_*n*_ is the corresponding singular value. Positive values in 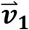 map to positive values in 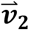, positive values in 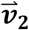 map to negative values in 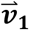, negative values in 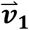 map to negative values in 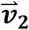 and negative values in 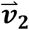 map to positive values in 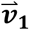:

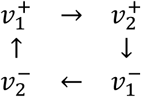

These are the four transformations we represent in **Figure 3D** and **Figure 7 – figure supplement 1**.

To determine whether transition modes were disrupted in mutants, we mapped mutant and control bouts into the behavioral space and computed the SVD of their transition frequency matrices. We compared the dot products of sibling control transition modes to transition modes obtained from the canonical dataset, and the dot products of mutant transition modes to sibling control transition modes (**Figure 7 – figure supplement 1**). We determined whether transition modes were significantly disrupted in mutants with a permutation test.

### Identification of behavioral modules

To identify behavioral modules, we combined information about bouts’ kinematics and transitions to generate a new kinematic-transition hybrid space. The kinematic nearest-neighbors graph was constructed from the DTW distances between exemplars as described above (*embedding postural dynamics in a behavioral space*). We constructed a transition space from the vector defining the second symmetric transition mode and the pair of vectors defining the first antisymmetric transition mode, and then calculated an orthogonal basis for this space. The first symmetric transition mode was excluded since it contains information about the prevalence of each kinematic motif in the data, rather than how motifs are chained together. Then, we warped the kinematic graph by multiplying the distances between adjacent nodes by the Euclidean distance between the corresponding exemplars in the transition space to generate our hybrid space. Then we proceeded with isomap embedding on this hybrid space, finding the shortest distance between each pair of motifs and taking the eigenvectors of the resulting matrix. This decomposition was dominated by two large eigenvalues, so we used a two-dimensional kinematic-transition space and performed hierarchical clustering using Ward’s method (**Figure 3E**). We set the threshold for separating clusters based on what we considered to provide the most parsimonious partitioning of bouts, referencing previously published literature and assessing whether further subdivision of the space produced interpretable clusters.

In **Figure 3F**, we colored points in the original behavioral space based on the cluster they were assigned in the hybrid kinematic-transition space. The transparency value in that graph was determined by the number of nearest neighbors that were assigned the same cluster label.

To produce average traces for the tail tip angle, we aligned all exemplars belonging to a given cluster using dynamic time warping and took the average of the aligned traces. The representative examples we show are those whose tail tip angle traces were most similar to each average.

### Modelling transitions between modules

For this analysis, we first identified every uninterrupted chain containing at least two bouts in our data which could be assigned a behavioral module, i.e. only chains of bouts from within a single recording trial (see *free-swimming behavioral assay*) and that could be embedded in the behavioral space (see *embedding postural dynamics in a behavioral space*). We then tested the ability of a series of Markov models – ranging from zeroth to fourth order – to predict each subsequent bout. For this purpose, we modelled each module as a state in a Markov process (allowing transitions to the same state, since fish can perform the same type of bout twice in a row). Each of our models contained seven states, *s*_1_, *s*_2_,…, *s*_7_, and we denote the current state, *X*_*t*_, the next state *X*_*t*+1_, the previous state *X*_*t*−1_, etc.

A zeroth order Markov model does not know the current state and therefore guesses the next state based simply on the distribution of bouts across all states:

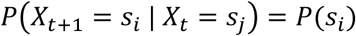

In a first order Markov model, the current state is known. To predict the next state, we considered all other times the current state was visited (*X*_*n*_) and observed which bout occurred next in the sequence:

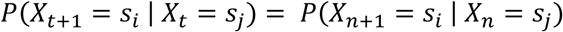

For the second-order Markov model, we took into account the last two states in a chain when predicting the next state:

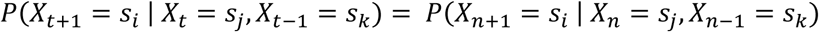

Similarly, for Markov models up to order, *m*, we predicted the next state:

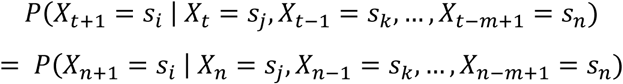

In **Figure 4B**, we show the probability that a given model predicts the next bout correctly starting from each behavioral module.

### Ethogram analysis

The ethogram in **Figure 4C,D** shows the first-order Markovian transition probabilities across all fish. To identify which transitions were significant, we used a permutation test. We shuffled the order of bouts *within* each fish 1000 times and recomputed the first-order Markovian transition probability matrices. This gave a distribution of transition probabilities between each pair of modules from which we could calculate the one-tailed p-values. We considered significant transitions as those that had a p-value < 0.05 after applying a Holm-Bonferroni correction (7^2^ = 49 comparisons).

To identify significantly altered transition in the mutants, we computed an ethogram for each fish and compared the distribution of transition probabilities across fish between groups with a series of Mann-Whitney U tests. We always compared mutants to sibling controls, and considered significant transitions as those that had a p-value < 0.05 after applying a Holm-Bonferroni correction.

### Spontaneous and prey capture chain analysis

To determine whether spontaneous and prey capture chains were disrupted in mutants, we computed the number of transitions within the groups {slow 2, burst, turn} and {J-turn, orientation, slow 1, capture strike} respectively, before a transition to a bout from the other group occurred. For each animal, we calculated the mean number of transitions within a group, and compared the distributions of these means between conditions, always comparing mutants to sibling controls, with a two-tailed Mann-Whitney U test (**Figure 7F,I**).

### Generating stimulus maps

To obtain stimulus maps for **Figure 4E** and **Figure 4 – figure supplement 1**, we took the first and last frames from the raw video data of each bout. We then performed background division (see *tail and eye tracking*) and binarized the images using a low threshold to remove pixel noise. We then aligned frames using the heading angle and swim bladder centroid obtained from the tracking. Next, we split bouts based on eye convergence state and behavioral module, and averaged the pre- and post-bout frames for each of these conditions. We then performed pixel-wise normalization of these average images using the mean and standard deviation of ∼90,000 frames randomly selected from periods when fish were not performing any bout. To obtain mirror-symmetric stimulus maps, we calculated the average of each image with its reflection.

We obtained the stimulus maps for **Figure 5F** and **Video 4** using the same process described above: background division, thresholding, alignment, averaging, normalization. To obtain the stimulus time series, we split captures into attack swims and S-strikes (see *capture strike analysis* below), and aligned videos in time to the onset of each bout.

To obtain stimulus maps for jaw movements (**Figure 6J**), we identified the onset of the bouts that immediately preceded each jaw movement. We calculated the average stimulus from the side for frames corresponding to these bout onsets as described above for the top view: background division, thresholding, binarization, alignment, averaging, normalization. We aligned frames using the centroid of the contour outlining the head and the pitch of the fish in the water. Normalization was performed with the average and standard deviation of ∼18,000 randomly selected frames.

### Capture strike analysis

For analysis relating to **Figure 5, Video 3** and **Video 4**, we defined capture strikes as bouts that were mapped to a kinematic motif that contains > 50% late prey capture bouts (**Figure 2D**). To determine the moment of capture in **Figure 5A**, we selected 100 random capture strikes and manually annotated the frames where the jaw was maximally extended.

For subsequent analysis, we only considered the 50 ms time window shown in **Figure 5A** (24-74 ms after the bout onset as determined by our bout segmentation algorithm) and proceeded with our general DTW-isomap embedding algorithm as described above (see *embedding postural dynamics in a behavioral space*). To generate the capture strike subspace, we computed the DTW distance between each pair of strikes, only allowing warping within a 6 ms (3 frames) time window. We used the resulting pairwise distance matrix directly for isomap embedding, keeping the first two dimensions. Note we did not perform an intermediate affinity propagation clustering for this dataset. We then performed KMeans clustering (sklearn.cluster.KMeans) with # clusters = 2 to classify strikes.

### Generating a behavioral space of jaw movements

To generate the jaw movement behavioral space in **Figure 6F**, we performed PCA on the jaw depression and cranial elevation traces across movement frames (see *jaw tracking*). We calculated the DTW distance (warping bandwidth = 10 ms) between each pair of movements projected onto the first principal component (**Figure 6E**), and performed isomap embedding using the resulting distance matrix. To identify clusters, we used <u>H</u>ierarchical <u>D</u>ensity-<u>B</u>ased <u>S</u>patial <u>C</u>lustering of <u>A</u>pplications with <u>N</u>oise (HDBSCAN) (hdbscan library, Python).

## Acknowledgements

The authors would like to thank all members of the Baier lab and colleagues at the Max Planck Institute of Neurobiology for discussions and helpful comments. In particular, we thank Thomas Helmbrecht, Johannes Larsch and Marco Dal Maschio for their constructive feedback on the manuscript and design of the analysis. We also thank Krasimir Slanchev for providing *lakritz* and *blumenkohl* larvae for mutant experiments. The schematic in Figure 7A was adapted with permission from unpublished work by Yvonne Kölsch. Funding was provided by the Max Planck Society.

## Author Contributions

Conceptualization, D.S.M., J.L.S. and H.B.; Methodology, D.S.M., J.C.D. and H.B.; Investigation, D.S.M.; Software, D.S.M. and J.C.D.; Formal Analysis, D.S.M.; Visualization, D.S.M.; Writing – Original Draft, D.S.M.; Writing – Review & Editing, J.L.S., J.C.D. and H.B.; Supervision, H.B.; Funding Acquisition, H.B.

## References

Anderson, D.J., and Perona, P. (2014). Toward a Science of Computational Ethology. Neuron 84, 18–31.

Berman, G.J., Choi, D.M., Bialek, W., and Shaevitz, J.W. (2014). Mapping the stereotyped behaviour of freely moving fruit flies. J. R. Soc. Interface 11, 20140672.

Berman, G.J., Bialek, W., and Shaevitz, J.W. (2016). Predictability and hierarchy in Drosophila behavior. Proc. Natl. Acad. Sci. 113, 11943–11948.

Bianco, I.H., and Engert, F. (2015). Visuomotor Transformations Underlying Hunting Behavior in Zebrafish. Curr. Biol. 25, 831–846.

Bianco, I.H., Kampff, A.R., and Engert, F. (2011). Prey Capture Behavior Evoked by Simple Visual Stimuli in Larval Zebrafish. Front. Syst. Neurosci. 5.

Bizzi, E., and Cheung, V.C. (2013). The neural origin of muscle synergies. Front. Comput. Neurosci. 7.

Borla, M.A., Palecek, B., Budick, S., and O’Malley, D.M. (2002). Prey Capture by Larval Zebrafish: Evidence for Fine Axial Motor Control. Brain. Behav. Evol. 60, 207–229.

Brown, A.E.X., and Bivort, B. de (2018). Ethology as a physical science. Nat. Phys. 1.

Brown, A.E.X., Yemini, E.I., Grundy, L.J., Jucikas, T., and Schafer, W.R. (2013). A dictionary of behavioral motifs reveals clusters of genes affecting Caenorhabditis elegans locomotion. Proc. Natl. Acad. Sci. 110, 791–796.

Budick, S.A., and O’Malley, D.M. (2000). Locomotor repertoire of the larval zebrafish: swimming, turning and prey capture. J. Exp. Biol. 203, 2565–2579.

Cande, J., Namiki, S., Qiu, J., Korff, W., Card, G.M., Shaevitz, J.W., Stern, D.L., and Berman, G.J. (2018). Optogenetic dissection of descending behavioral control in Drosophila. ELife 7, e34275.

Coen, P., Clemens, J., Weinstein, A.J., Pacheco, D.A., Deng, Y., and Murthy, M. (2014). Dynamic sensory cues shape song structure in Drosophila. Nature 507, 233–237.

Del Vecchio, D., Murray, R.M., and Perona, P. (2003). Decomposition of human motion into dynamics-based primitives with application to drawing tasks. Automatica 39, 2085–2098.

Egnor, S.E.R., and Branson, K. (2016). Computational Analysis of Behavior.

Ewert, J.-P. (1987). Neuroethology of releasing mechanisms: Prey-catching in toads. Behav. Brain Sci. 10, 337–368.

Flash, T., and Hochner, B. (2005). Motor primitives in vertebrates and invertebrates. Curr. Opin. Neurobiol. 15, 660–666.

Frey, B.J., and Dueck, D. (2007). Clustering by Passing Messages Between Data Points. Science 315, 972–976.

Gahtan, E., Tanger, P., and Baier, H. (2005). Visual Prey Capture in Larval Zebrafish Is Controlled by Identified Reticulospinal Neurons Downstream of the Tectum. J. Neurosci. 25, 9294–9303.

Girdhar, K., Gruebele, M., and Chemla, Y.R. (2015). The Behavioral Space of Zebrafish Locomotion and Its Neural Network Analog. PLOS ONE 10, e0128668.

Haesemeyer, M., Robson, D.N., Li, J.M., Schier, A.F., and Engert, F. (2015). The Structure and Timescales of Heat Perception in Larval Zebrafish. Cell Syst. 1, 338–348.

Helmbrecht, T.O., dal Maschio, M., Donovan, J.C., Koutsouli, S., and Baier, H. (2018). Topography of a Visuomotor Transformation. Neuron 100, 1429–1445.e4.

Hernández, L.P., Barresi, M.J.F., and Devoto, S.H. (2002). Functional Morphology and Developmental Biology of Zebrafish: Reciprocal Illumination from an Unlikely Couple. Integr. Comp. Biol. 42, 222–231.

Horstick, E.J., Bayleyen, Y., Sinclair, J.L., and Burgess, H.A. (2017). Search strategy is regulated by somatostatin signaling and deep brain photoreceptors in zebrafish. BMC Biol. 15, 4.

Jouary, A., and Sumbre, G. (2016). Automatic classification of behavior in zebrafish larvae. BioRxiv 052324.

Kato, S., Kaplan, H.S., Schrödel, T., Skora, S., Lindsay, T.H., Yemini, E., Lockery, S., and Zimmer, M. (2015). Global Brain Dynamics Embed the Motor Command Sequence of Caenorhabditis elegans. Cell 163, 656–669.

Katsov, A.Y., Freifeld, L., Horowitz, M., Kuehn, S., and Clandinin, T.R. (2017). Dynamic structure of locomotor behavior in walking fruit flies. ELife 6, e26410.

Kay, J.N., Finger-Baier, K.C., Roeser, T., Staub, W., and Baier, H. (2001). Retinal Ganglion Cell Genesis Requires lakritz, a Zebrafish atonal Homolog. Neuron 30, 725–736.

Krakauer, J.W., Ghazanfar, A.A., Gomez-Marin, A., MacIver, M.A., and Poeppel, D. (2017). Neuroscience Needs Behavior: Correcting a Reductionist Bias. Neuron 93, 480–490.

Larsch, J., and Baier, H. (2018). Biological Motion as an Innate Perceptual Mechanism Driving Social Affiliation. Curr. Biol. 28, 3523–3532.e4.

Long, M.A., Jin, D.Z., and Fee, M.S. (2010). Support for a synaptic chain model of neuronal sequence generation. Nature 468, 394–399.

Marques, J.C., Lackner, S., Félix, R., and Orger, M.B. (2018). Structure of the Zebrafish Locomotor Repertoire Revealed with Unsupervised Behavioral Clustering. Curr. Biol. 28, 181–195.e5.

McClenahan, P., Troup, M., and Scott, E.K. (2012). Fin-Tail Coordination during Escape and Predatory Behavior in Larval Zebrafish. PLOS ONE 7, e32295.

McElligott, M.B., and O’Malley, D.M. (2005). Prey Tracking by Larval Zebrafish: Axial Kinematics and Visual Control. Brain. Behav. Evol. 66, 177–196.

Neuhauss, S.C.F., Biehlmaier, O., Seeliger, M.W., Das, T., Kohler, K., Harris, W.A., and Baier, H. (1999). Genetic Disorders of Vision Revealed by a Behavioral Screen of 400 Essential Loci in Zebrafish. J. Neurosci. 19, 8603–8615.

Patterson, B.W., Abraham, A.O., MacIver, M.A., and McLean, D.L. (2013). Visually guided gradation of prey capture movements in larval zebrafish. J. Exp. Biol. 216, 3071–3083.

Sakoe, H., and Chiba, S. (1978). Dynamic programming algorithm optimization for spoken word recognition. IEEE Trans. Acoust. Speech Signal Process. 26, 43–49.

Seeds, A.M., Ravbar, P., Chung, P., Hampel, S., Jr, F.M.M., Mensh, B.D., and Simpson, J.H. (2014). A suppression hierarchy among competing motor programs drives sequential grooming in Drosophila. ELife 3, e02951.

Semmelhack, J.L., Donovan, J.C., Thiele, T.R., Kuehn, E., Laurell, E., and Baier, H. (2014). A dedicated visual pathway for prey detection in larval zebrafish. ELife 3, e04878.

Smear, M.C., Tao, H.W., Staub, W., Orger, M.B., Gosse, N.J., Liu, Y., Takahashi, K., Poo, M., and Baier, H. (2007). Vesicular Glutamate Transport at a Central Synapse Limits the Acuity of Visual Perception in Zebrafish. Neuron 53, 65–77.

Stephens, G.J., Johnson-Kerner, B., Bialek, W., and Ryu, W.S. (2008). Dimensionality and Dynamics in the Behavior of C. elegans. PLOS Comput. Biol. 4, e1000028.

Szigeti, B., Deogade, A., and Webb, B. (2015). Searching for motifs in the behaviour of larval Drosophila melanogaster and Caenorhabditis elegans reveals continuity between behavioural states. J. R. Soc. Interface 12, 20150899.

Tao, L., Ozarkar, S., Beck, J.M., and Bhandawat, V. (2019). Statistical structure of locomotion and its modulation by odors. ELife 8, e41235.

Tenenbaum, J.B., Silva, V. de, and Langford, J.C. (2000). A Global Geometric Framework for Nonlinear Dimensionality Reduction. Science 290, 2319–2323.

Tinbergen, N. (1951). The Study of Instinct (Oxford: Oxford University Press).

Trivedi, C.A., and Bollmann, J.H. (2013). Visually driven chaining of elementary swim patterns into a goal-directed motor sequence: a virtual reality study of zebrafish prey capture. Front. Neural Circuits 7.

Vogelstein, J.T., Park, Y., Ohyama, T., Kerr, R.A., Truman, J.W., Priebe, C.E., and Zlatic, M. (2014). Discovery of Brainwide Neural-Behavioral Maps via Multiscale Unsupervised Structure Learning. Science 344, 386–392.

Wiltschko, A.B., Johnson, M.J., Iurilli, G., Peterson, R.E., Katon, J.M., Pashkovski, S.L., Abraira, V.E., Adams, R.P., and Datta, S.R. (2015). Mapping Sub-Second Structure in Mouse Behavior. Neuron 88, 1121–1135.

Zimmermann, M.J.Y., Nevala, N.E., Yoshimatsu, T., Osorio, D., Nilsson, D.-E., Berens, P., and Baden, T. (2018). Zebrafish Differentially Process Color across Visual Space to Match Natural Scenes. Curr. Biol. 28, 2018–2032.e5.

