## Supplemental Figures for "Deconstructing hunting behavior reveals a tightly coupled stimulus-response loop"

**Video 1.** Free-swimming behavioral tracking. Example six seconds of behavioral data (same data as **Figure 1B,C**), containing a hunting sequence. We track the animal in each frame (left), and extract the kinematics of its movements (right). Tracking is performed offline. We track the angle of the left and right eyes (top, green and magenta traces respectively) and the angle of the tail at 50 equally-spaced points (middle). Zebrafish larvae swim in bouts, which we detect automatically using the tail tip angle (bottom). Behavior is recorded at 500 fps. Playback is 50 fps (one tenth actual speed).

**Video 2.** Postural dynamics of the tail are a trajectory in three-dimensional principal component space. Same recording as **Figure 1B,C** and **Video 1**. Top right: head-stabilized fish from raw video data. Left: in each frame, the tail shape is approximated by a point in three-dimensional postural space, defined by the principal components. The shape of the tail changes as the fish swims and traces a trajectory in the postural space. Bottom right: estimated pose dynamics reconstructed from the three-dimensional trajectory through postural space.

**Video 3.** Examples of attack swims and S-strikes. Nine representative examples each of attack swims and S-strikes. Examples are from a mix of animals and are aligned to the bout onset. Note, only the first 100 ms of the bouts are shown, even though some may continue for longer. The attempted capture event (when the jaw opens) occurs after approximately 60 ms. Note the prominent and highly stereotyped S-bend of the tail during the S-strikes. Behavior is recorded at 500 fps. Playback is 5 fps (one hundredth actual speed).

**Video 4.** Evolution of the stimulus during prey capture. Normalized average stimulus evolution for hunting sequences the lead to either an attack swim (left) or an S-strike (middle). Brighter colors signify a more stereotyped (i.e. higher probability density) prey position in that part of the image at a given time. Hunting sequences are aligned to the onset of the capture strike (0 ms). Images are normalized using the mean and standard deviation from approximately 90,000 randomly selected frames. White contour shows outline of the fish. Changes in density either side of the fish contour signify fin movements. Right: maximum prey density along axial and horizontal axes in the anterior visual field.

**Video 5.** Free-swimming behavioral tracking from two angles simultaneously. Example four seconds of behavioral data (same data as **Figure 6B,C**), containing two hunting sequences. We track the animal in each frame (left), and extract the kinematics of its movements (right). Tracking is performed offline. We track the depression of the jaw (left image: length of cyan bar; right: top black trace) and elevation of the cranium (left image: angles between red lines; right top gray trace). To perform tail tracking, we measure the angle of the tail at 50 equally-spaced points and detect bouts automatically (right middle). We also tracking the pitch of the fish in the water (left image: angle of longer red line relative to horizontal; right: bottom black trace). Capture events occur when cranial elevation spikes. Note the rising pitch of the fish prior to these events. Behavior is recorded at 400 fps. Playback is 40 fps (one tenth actual speed).

**Video 6.** High-resolution recording of the capture strike jaw movement. We used two cameras to record a zebrafish larva hunting a paramecium (indicated by the black circle at the beginning of the recording) simultaneously from above and from the side at higher spatial resolution. Note the extreme jaw depression and cranial elevation achieved by the fish during prey capture. Time zero indicates moment the prey is consumed, when jaw depression and cranial elevation are at a maximum. Behavior is recorded at 191 fps. Playback is 6 fps (approximately 30x slower than actual speed).

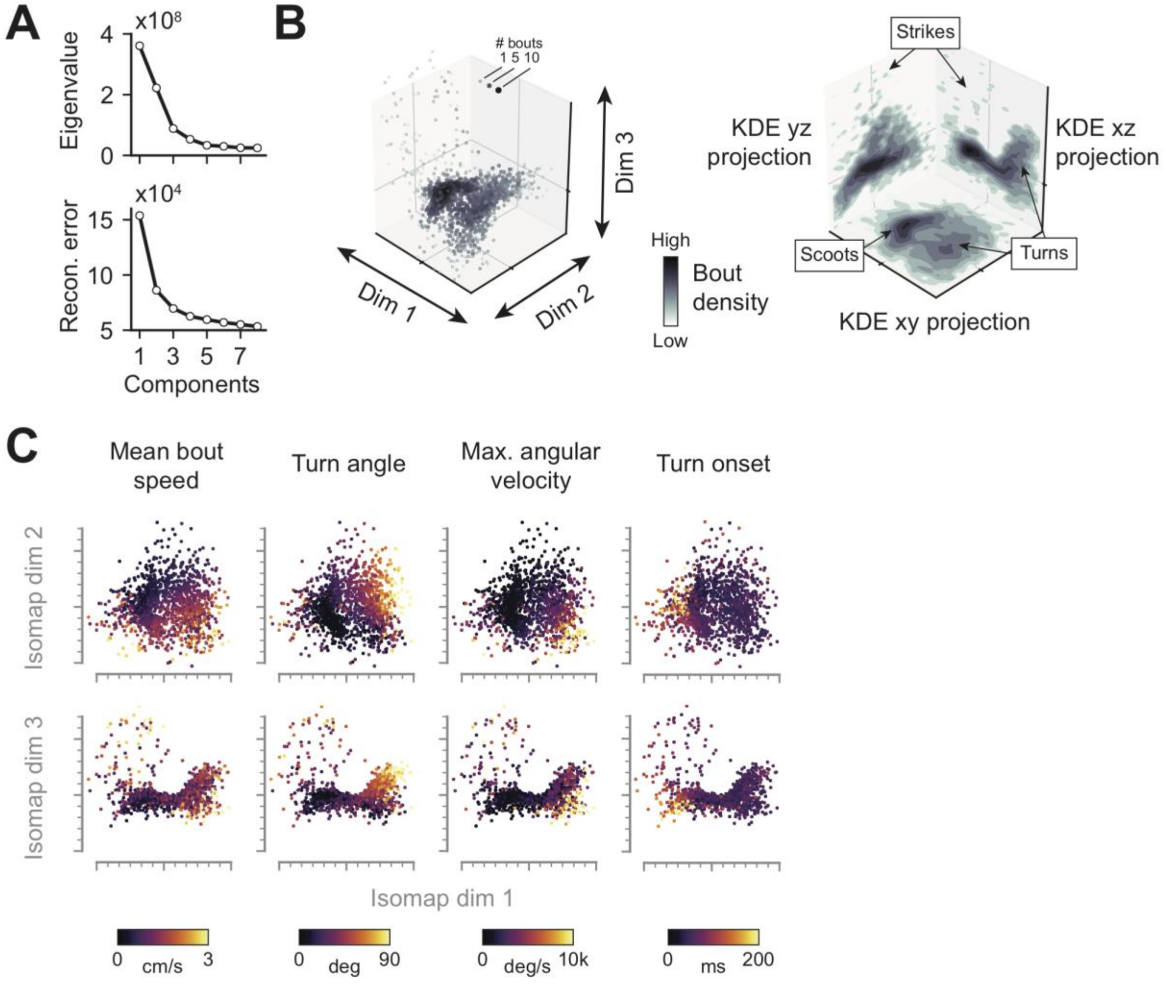


**Figure 2 – figure supplement 1.** Structure of zebrafish behavioral space. (**A**) Eigenvalues and reconstruction errors corresponding to the isomap embedding used to generate the behavioral space. (**B**) Density of bouts within the behavioral space. Left: bouts represented in full 3D space. Right: Kernel density estimation (KDE) of bouts projected onto planes defined by each pair of axes in the behavioral space. (**C**) Mean speed, turn angle, maximum angular velocity and time of turn onset of each exemplar in the behavioral space. Values represent median across all bouts mapped to each exemplar.

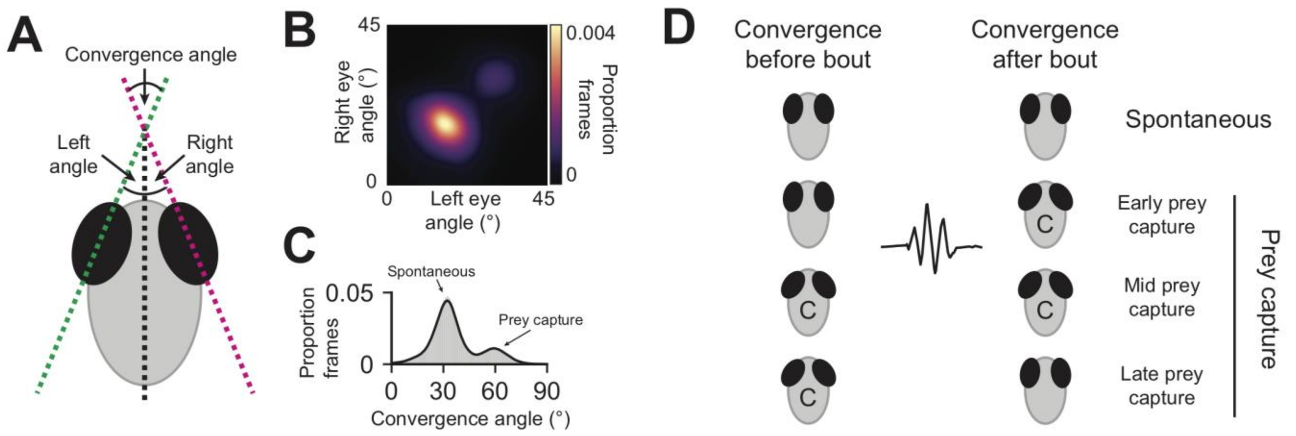


**Figure 2 – figure supplement 2.** Prey capture and spontaneous swimming are defined by eye convergence. (**A**) Schematic showing calculation of left eye, right eye, and eye convergence angles. (**B**) 2D histogram of left and right eye angles across all frames of all recordings. (**C**) Bimodal distribution of eye convergence angles across all frames of all recordings. Such distributions are calculated for each fish individually and the eye convergence threshold is defined by the antimode of this distribution. (**D**) Schematic showing classification of bouts into spontaneous, early prey capture, mid prey capture, or late prey capture phases. The latter three are all considered as prey capture.

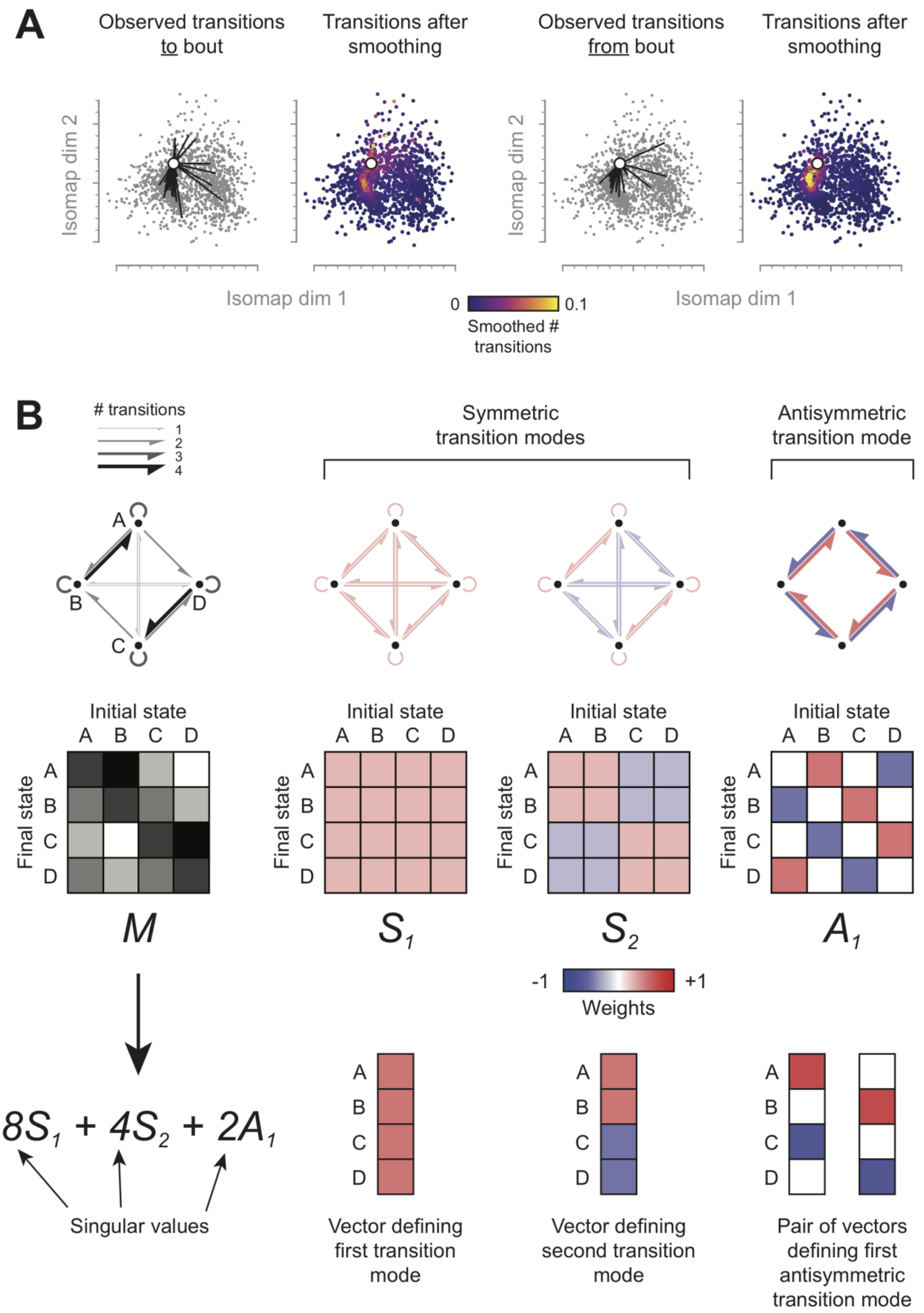
A
Observed transitions to bout
Transitions after smoothing
Observed transitions from bout
Transitions after smoothing
Isomap dim 2
Isomap dim 1
0
0.1
Smoothed # transitions
B
### transitions
1
2
3
4
Symmetric transition modes
Antisymmetric transition mode
A
B
C
D
Initial state
Final state
M
S1
S2
A1
-1
+1
Weights
8S1 + 4S2 + 2A1
Singular values
Vector defining first transition mode
Vector defining second transition mode
Pair of vectors defining first antisymmetric transition mode

**Figure 3 – figure supplement 1.** Explanation of singular value decomposition. (**A**) Smoothing transitions in the behavioral space. To facilitate identification of transition modes by singular-value decomposition, transitions to and from each bout are smoothed in the behavioral space. Note, that only the first two dimensions of the behavioral space are shown, although transitions are smoothed in all three dimensions. See also Methods. (**B**) Explanation of SVD using a hypothetical example containing only four behavioral states. Top left: number of transitions observed between each pair of states, lettered A to D counterclockwise starting at the top, depicted as both an ethogram and a transition frequency matrix, *M*. Top right: transition modes identified through singular-value decomposition of *M*, depicted as ethograms and matrices. Note that transition modes allow for “negative” transitions as well as positive ones. Bottom right: each symmetric transition mode is associated with a vector, and the antisymmetric transition mode is associated with a pair of orthogonal vectors. In **Figure 3C,D**, bouts with the same sign in a singular vector are depicted together. Bottom left: the complete transition frequency matrix, *M*, can be reconstituted by summing the transition modes multiplied by their corresponding singular values. See also Methods.

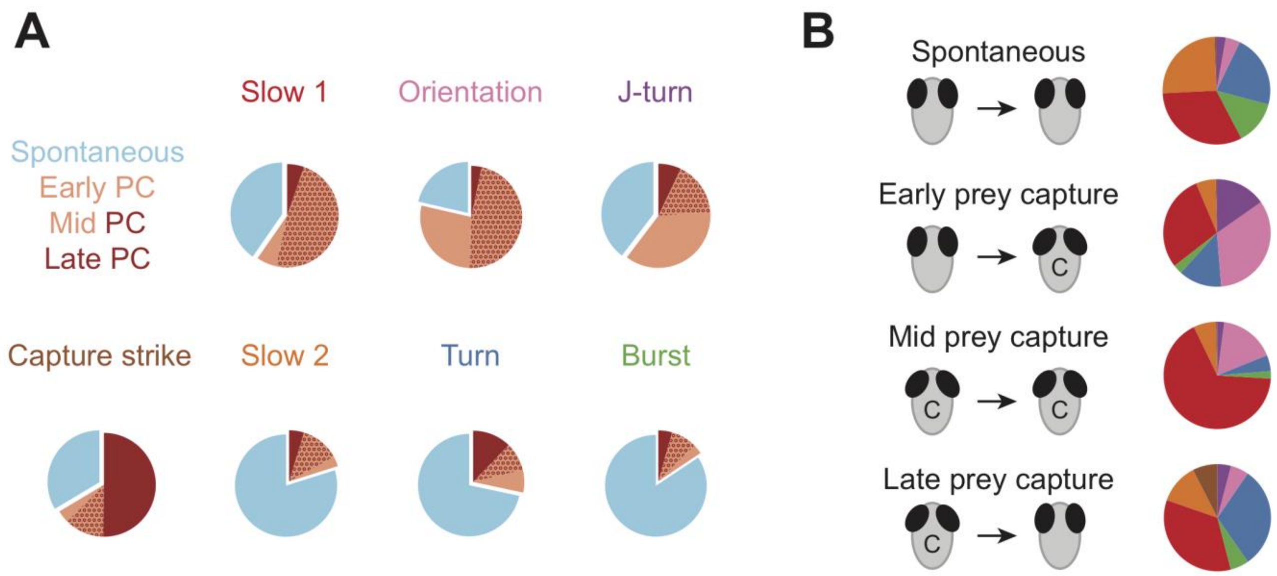


**Figure 3 – figure supplement 2.** Behavioral modules and eye convergence. (**A**) Proportion of bouts in each module that occur during spontaneous swimming (blue), early prey capture (pink), mid prey capture (pink/red), or late prey capture (red). Prey capture modules are defined as those that predominantly occur during eye convergence. (**B**) Proportion of bouts during each eye convergence phase belonging to each behavioral module.

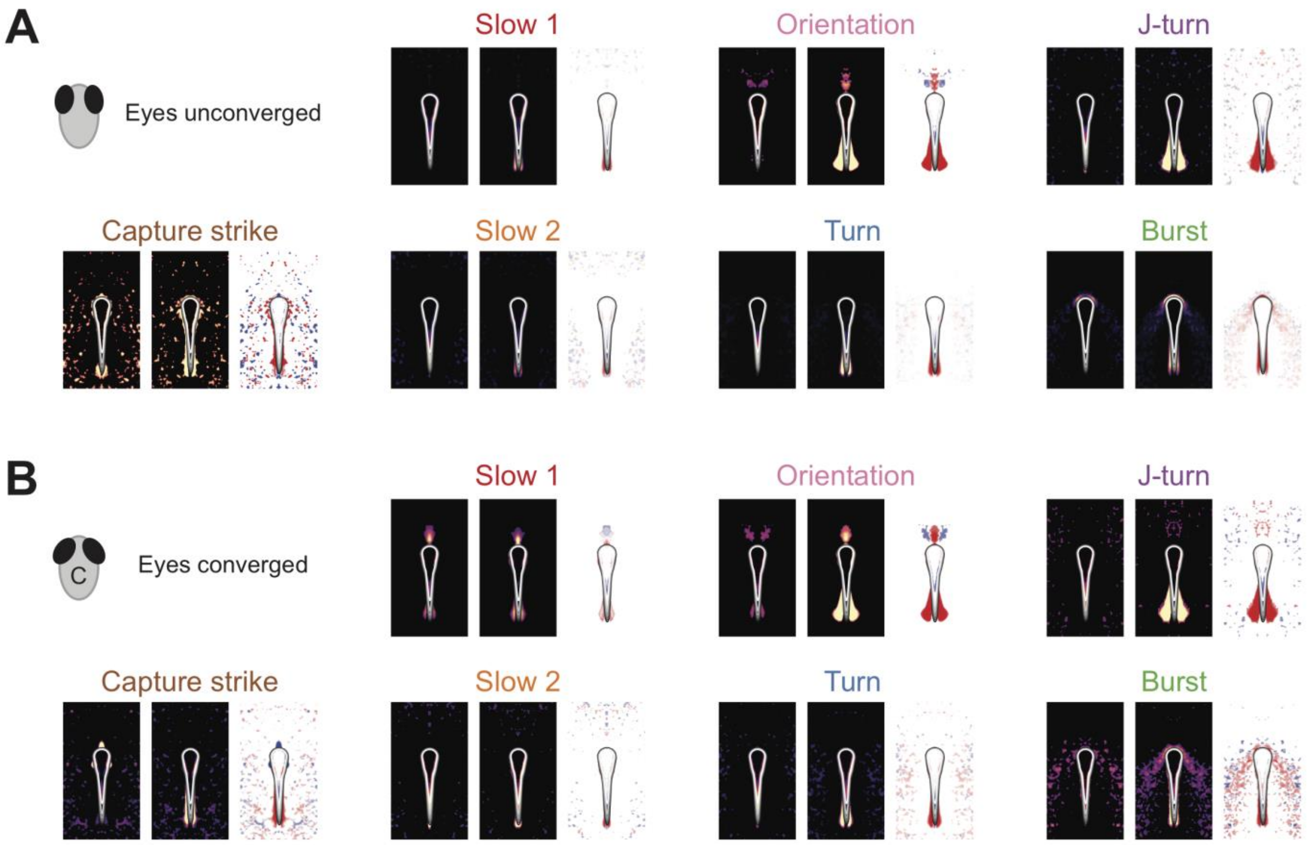


**Figure 4 – figure supplement 1.** Stimulus maps for modules during spontaneous swimming and prey capture. (**A**) Average stimulus before and after each type of bout when eyes are not converged (spontaneous swimming). (**B**) Average stimulus before and after each type of bout when eye are converged (early, mid and late prey capture). Maps show average pixel intensity around fish across all bouts before (left) and after (middle) each module, expressed as a z-score (normalized using mean and standard deviation of 90,000 randomly selected frames). Difference is shown on the right. Images are thresholded using 95$^{th}$ percentile. Contour shows outline of the fish.

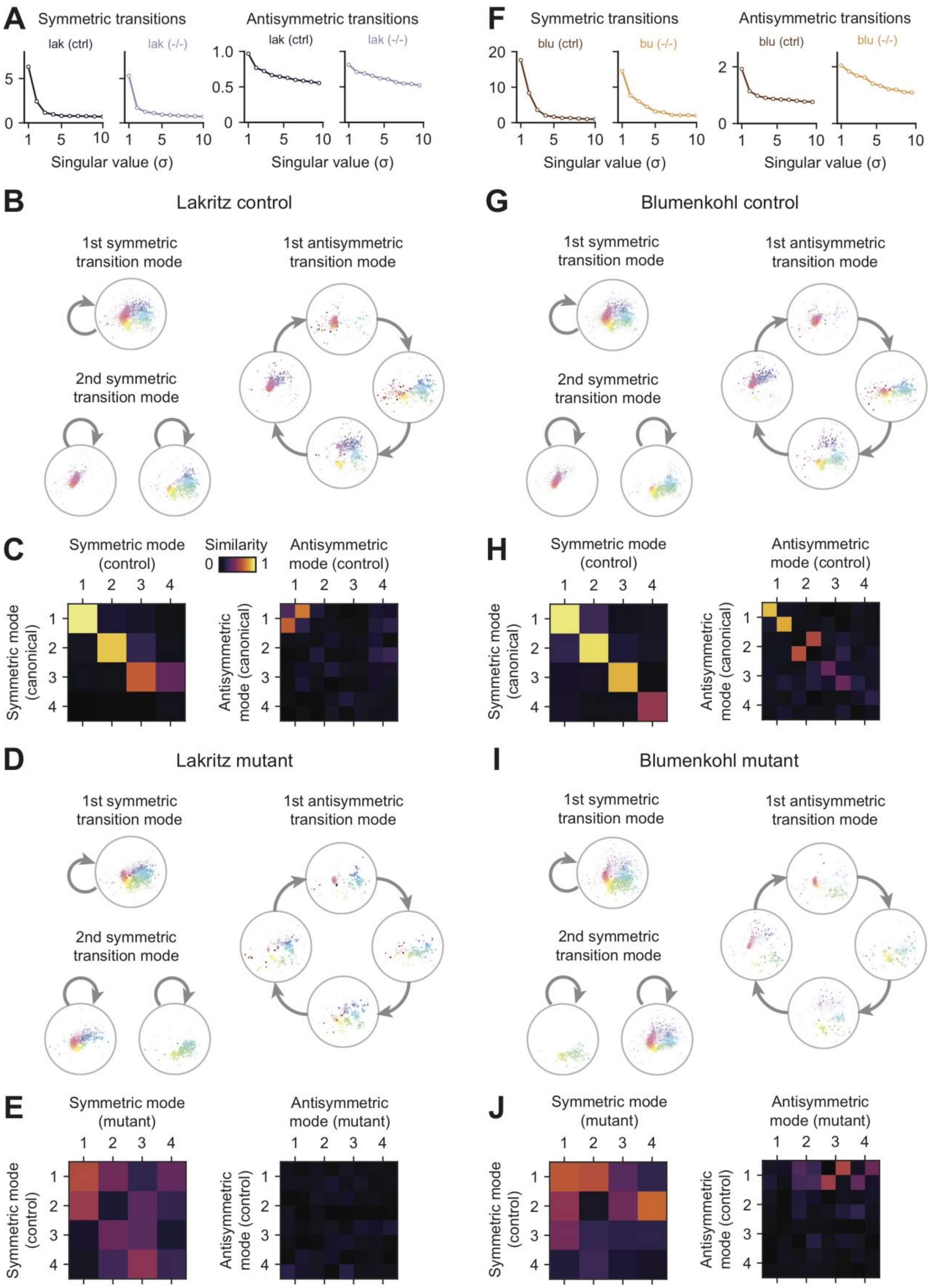
A
Symmetric transitions
lak (ctrl)
lak (-/-)
Antisymmetric transitions
lak (ctrl)
lak (-/-)
Singular value (σ)
Singular value (σ)
B
Lakritz control
1st symmetric transition mode
2nd symmetric transition mode
1st antisymmetric transition mode
C
Symmetric mode (control)
Similarity
Antisymmetric mode (control)
Symmetric mode (canonical)
Antisymmetric mode (canonical)
D
Lakritz mutant
1st symmetric transition mode
2nd symmetric transition mode
1st antisymmetric transition mode
E
Symmetric mode (mutant)
Antisymmetric mode (mutant)
Symmetric mode (control)
Antisymmetric mode (control)
F
Symmetric transitions
blu (ctrl)
bu (-/-)
Antisymmetric transitions
blu (ctrl)
blu (-/-)
Singular value (σ)
Singular value (σ)
G
Blumenkohl control
1st symmetric transition mode
2nd symmetric transition mode
1st antisymmetric transition mode
H
Symmetric mode (control)
Antisymmetric mode (control)
Symmetric mode (canonical)
Antisymmetric mode (canonical)
I
Blumenkohl mutant
1st symmetric transition mode
2nd symmetric transition mode
1st antisymmetric transition mode
J
Symmetric mode (mutant)
Antisymmetric mode (mutant)
Symmetric mode (control)
Antisymmetric mode (control)

**Figure 7 – figure supplement 1.** Transition modes of *lakritz* and *blumenkohl* mutants. (**A**) Singular values of the transition matrices obtained from *lak* controls and mutants. (**B**) First two symmetric and first antisymmetric transition modes of *lak* controls. (**C**) Comparison of *lak* control transition modes to those of the canonical dataset (**Figure 3**). (**D**) First two symmetric and first antisymmetric transition modes of *lak* mutants. (**E**) Comparison of *lak* mutant transition modes to *lak* control transition modes. (**F**) Singular values of the transition matrices obtained from *blu* controls and mutants. (**G**) First two symmetric and first antisymmetric transition modes of *blu* controls. (**H**) Comparison of *blu* control transition modes to those of the canonical dataset (**Figure 3**). (**I**) First two symmetric and first antisymmetric transition modes of *blu* mutants. (**J**) Comparison of *blu* mutant transition modes to *blu* control transition modes. In (**C**), (**E**), (**H**), and (**J**) similarity is defined as the absolute dot product between vectors representing transition modes. Note that, for antisymmetric transitions, modes between datasets may be related by a 90° rotation.
